# Systematic cell-type resolved transcriptomes of 8 tissues in 8 lab and wild-derived mouse strains captures global and local expression variation

**DOI:** 10.1101/2025.04.21.649844

**Authors:** Elisabeth Rebboah, Ryan Weber, Elnaz Abdollahzadeh, Nikhila Swarna, Delaney K. Sullivan, Diane Trout, Fairlie Reese, Heidi Yahan Liang, Ghassan Filimban, Parvin Mahdipoor, Margaret Duffield, Romina Mojaverzargar, Erisa Taghizadeh, Negar Fattahi, Negar Mojgani, Haoran Zhang, Rebekah K. Loving, Maria Carilli, A. Sina Booeshaghi, Shimako Kawauchi, Ingileif B. Hallgrímsdóttir, Brian A. Williams, Grant R. MacGregor, Lior Pachter, Barbara J. Wold, Ali Mortazavi

**Affiliations:** Developmental and Cell Biology, University of California Irvine, Irvine, USA; Center for Complex Biological Systems, University of California Irvine, Irvine, USA; Division of Biology and Biological Engineering, California Institute of Technology, Pasadena, USA; David Geffen School of Medicine, University of California Los Angeles, Los Angeles, USA; Department of Bioengineering, University of California Berkeley, Berkeley, USA; Transgenic Mouse Facility, University of California Irvine, Irvine, USA

## Abstract

Mapping the impact of genomic variation on gene expression facilitates an understanding of the molecular basis of complex phenotypic traits and disease predisposition. Mouse models provide a controlled and reproducible framework for capturing the breadth of genomic variation observed in different genotypes across a wide variety of tissues. As part of the IGVF consortium’s effort to catalog the effects of genetic variation, we uniformly characterized the transcriptomes of eight tissues from each mouse founder strain used to derive the Collaborative Cross strains, comprising five classical laboratory inbred strains and three wild-derived inbred strains. We sequenced samples from four male and four female replicates per tissue using single-nucleus RNA-seq to generate an “8-cube” dataset of 5.2 million nuclei across 106 cell types and cell states. As expected, the overall extent of transcriptome variation correlates positively with genetic divergence across the strains with the greatest differential between PWK/PhJ and CAST/EiJ. At the individual tissue level, heart and brain are relatively more similar across strains compared with gonads, adrenal, skeletal muscle, kidney, and liver. Further analyses revealed substantial strain variation, often concentrated in a few cell types as well as cell-state signatures that especially reflect strain-associated immune and metabolic trait differences. The founder 8-cube dataset provides rich transcriptome variation signatures to help explain strain-specific phenotypic traits and disease states, as illustrated by examples in tissue-resident immune cells, muscle degeneration, kidney sex differences, and the hypothalamicpituitary-adrenal axis. This data further provides a systematic foundation for the analysis of these tissues in the founder strains as well as the Collaborative Cross.

## Introduction

Understanding how genetic variation influences gene expression is fundamental to deciphering the molecular mechanisms that underlie complex traits, diseases, and developmental processes ^1,2^. It also promises new inroads into evolution of these traits. Genetic variants, including single nucleotide polymorphisms (SNPs) and structural variants, alter gene expression by modifying regulatory elements such as promoters, enhancers, and transcription factor binding sites or by creating local uncompensated gene dosage differences. These changes contribute to phenotypic variation among individuals, including differences in disease susceptibility. Investigating the effects of genetic variation on gene expression across tissues and at the cellular level promises insight into the regulatory networks that control cellular function. Because cell-type-specific gene expression patterns are essential for proper tissue and system function, quantifying the impact of genetic variants on these patterns is expected to reveal regulatory mechanisms and potential therapeutic targets. A central goal of the IGVF (Impact of Genomic Variation on Function) consortium is to comprehensively map the effects of genetic variation on molecular traits, including gene expression, across multiple tissues and disease contexts in both humans and mice at single-cell resolution ^3^.

Recombinant inbred mouse strains provide a powerful platform for investigating complex mammalian traits and diseases ^4–6^. These strains are generated by crossing genetically diverse founders to produce F1 hybrids, followed by intercrossing to generate F2 offspring, which are then inbred for at least 20 generations ^4^. This process yields stable recombinant inbred strains with near-homozygous genomes that are genetically identical within each strain but distinct between strains, making them ideal for genetic mapping studies ^4^. A caveat is that this process eliminates genotypes with substantially disadvantageous homozygous phenotypes. More recently, eight diverse founder strains were combined using an eight-way funnel breeding scheme to produce the widely-used Collaborative Cross (CC) recombinant inbred panel for studying complex traits ^7^. The CC panel is intended to capture and to model natural genetic variation while preserving the reproducibility and consistency afforded by inbred strains. Each CC strain represents a unique combination of haplotypes from eight founder strains, comprising five laboratory strains (C57BL/6J, NOD/ShiLtJ, NZO/HlLtJ, A/J, 129S1/SvImJ) and three wild-derived strains (WSB/J, PWK/PhJ, and CAST/EiJ). Together, these eight founders capture over 90% of the genetic variation available in *Mus musculus*, enabling the study of a broad spectrum of traits within a reproducible, controlled, and genetically defined framework ^8–12^. The CC panel has become a cornerstone of mouse systems genetics, enabling studies across a wide range of fields including infectious disease ^13–18^, cancer ^12,19,20^, metabolism and metabolic disease ^21,22^, neurobiology and behavior ^23,24^, toxicology and pharmacogenetics ^25–30^, and the gut microbiome ^31–33^. Despite the widespread use of the CC panel in dozens of studies, comprehensive transcriptomic data across the eight founder strains, particularly at single-cell resolution, has remained limited.

While C57BL/6J (“B6”) is the most widely-used laboratory strain, each of the other seven CC founder strains have also been leveraged for specific areas of disease research. For example, NOD/ShiLtJ (non-obese diabetic, “NOD”) is a well-established model for type 1 diabetes, while NZO/HlLtJ (New Zealand obese, “NZO”) is used to study type 2 diabetes ^34,35^. A/J (“AJ”) serves as a model for asthma ^36^, emphysema ^37^, and age-onset muscular dystrophy ^38^. Some strains are also preferred for particular experimental approaches. For example, 129S1/SvImJ (129S1) was historically a preferred background for deriving embryonic stem cells (ESCs), leading to the production of widely used cell lines such as CJ7^39^. ESC lines are now available for all eight founders ^40^. In contrast to the classical laboratory strains, each wild-derived strain originated from individuals trapped from a wild mouse population that were intercrossed until inbred. The wild-derived founders capture genetic variation from natural populations and represent the three main *Mus musculus* subspecies that diverged approximately one million years ago ^41^. These subspecies are *Mus musculus domesticus* (represented by WSB/J, “WSB”), *Mus musculus musculus* (represented by PWK/PhJ, “PWK”), and *Mus musculus castaneus* (represented by CAST/EiJ, “CAST”). These wild-derived strains also exhibit distinct phenotypic traits. For example, CAST mice can mount a significant regenerative response following neuronal injury in their central nervous system ^42^. CAST are also immune to flaviviruses but highly susceptible to other viruses such as orthopoxviruses, COVID-19, and influenza A, while PWK show resistance to influenza A ^43–46^. WSB mice, despite typical fertility rates, have significantly reduced sperm count and altered sperm morphology ^47^. Collectively, the founder strains encompass approximately 23 million unique SNPs and 350 million base pairs of structural variation compared to B6, providing a rich resource for studying how genetic variation shapes phenotypes ^40,48^. By capturing cell-type-specific gene expression across a broad range of biological systems, this dataset offers a reference for interpreting gene expression data from past and future CC-based studies.

Here, we used single-nucleus RNA-seq (snRNA-seq) to characterize gene expression in the eight founder strains of the Collaborative Cross panel across eight distinct tissue groups: (1) cortex and hippocampus, (2) diencephalon and pituitary gland, (3) muscle, (4) heart, (5) liver, (6) kidney, (7) adrenal gland, and (8) male and female gonads. We sequenced four 10-week-old adult males and females per genotype and recovered 106 heterogeneous cell types and states. By doing so, we uncovered strain-specific transcriptional differences across tissues that are correlated to variation in specific cell states.

## Results

### Experimental design and quality control of the 8-cube dataset

We collected eight tissues or tissue groups across eight genetically diverse founder strains from young adult mice at 10 weeks of age, sampling at least four males and four females per strain. Whenever possible, we profiled the same set of 64 individuals across all tissues, resulting in a coordinated whole-body dataset for each mouse. Our core dataset thus consists of 8 tissues by 8 strains by 8 replicates, for a total of 512 samples which we call the “8-cube” dataset (Fig. 1a). During sample collection, we recorded detailed metadata, including body and tissue weight, collection time, and estrus stage in females (Supp. Table 1). As expected from prior studies, NZO was the heaviest strain with an average of 43 grams, i.e. 1.95 times more than the average weight of 22 grams across all strains (Supp. Fig. 1a-i). By contrast, the wild-derived strains were typically the lightest with a mean weight of 14.4 grams. Sexual dimorphism was apparent in both body weight (with males weighing 16% more than females on average within each strain) as well as in certain tissues such as muscle and kidney, where male tissues weigh 31.1% and 20.5% more than female tissues, respectively. B6 and NOD had the largest sexual dimorphism in overall body weight, with males weighing almost 1.5 times more than females. We found strong positive correlation between body weight and tissue weights for heart, muscle, male gonads (testes and epididymis), kidney, and liver (R^2^ > 0.5, Supp. Fig. 1e-i), whereas adrenal glands and brain regions showed no correlation (R^2^ < 0.05, Supp. Fig. 1a-c). Female gonads, which include ovaries and oviducts, are relatively small and may be more affected by measurement limitations, showed moderate correlation with body weight (R^2^ = 0.43, Supp. Fig. 1d). Prior studies support these general trends. In 6 to 8 week old rats, body weight shows significant linear relationships with liver, kidneys, heart, and testes weights, along with weaker but still significant associations with adrenal and brain weights ^49^. In mice, brain weight increases in proportion to body weight during early postnatal development but stabilizes around postnatal day 15, even as body weight continues to rise ^50^, consistent with our observation of no correlation between body weight and brain region weight in 10-week-old animals. Together, these findings validate our trends and emphasize that while body weight remains a relevant biological variable, its relationship to tissue weight is context-dependent in adult mice.

**Figure 1.**
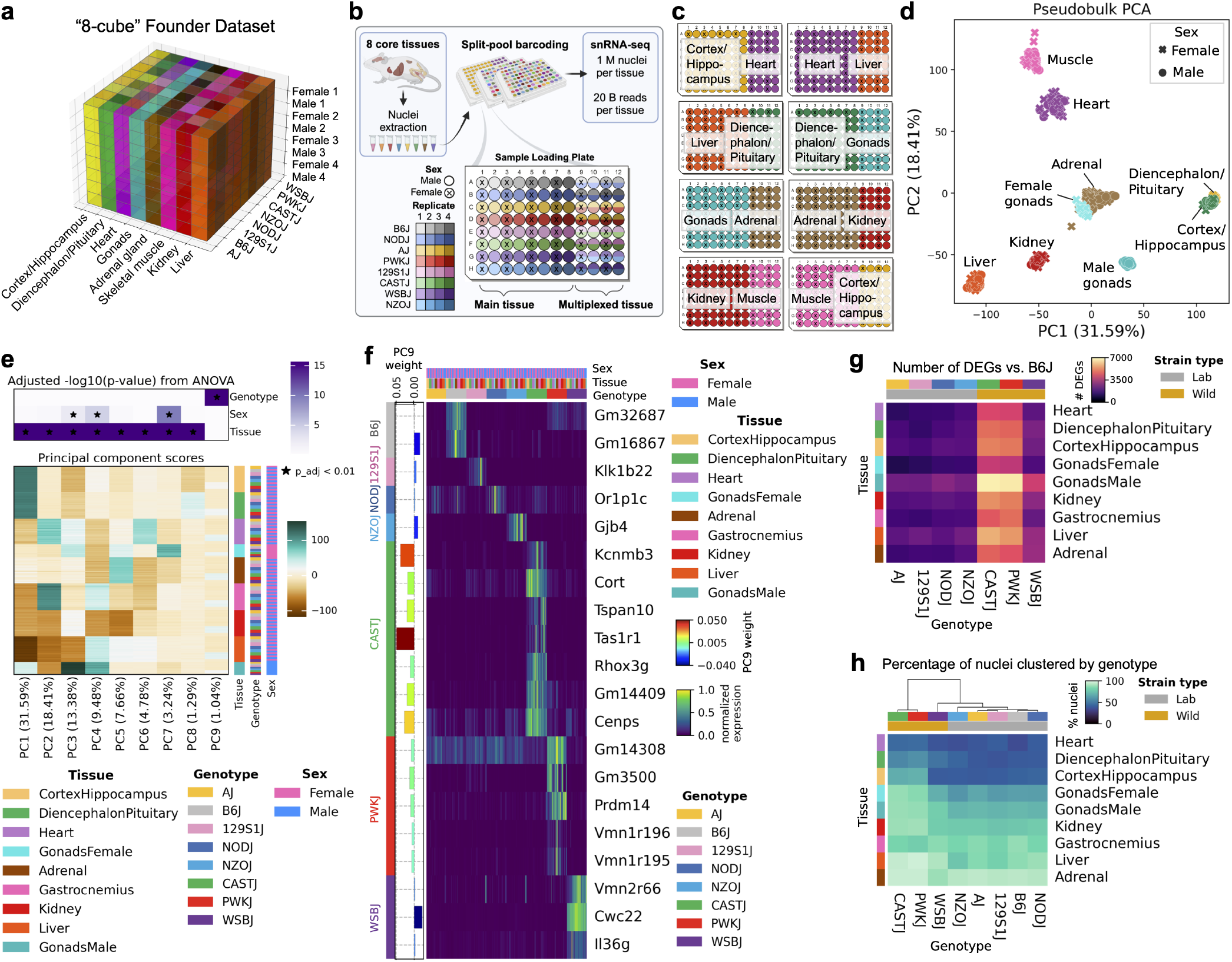
Overview of the IGVF mouse dataset in 8 founder genotypes. **a**, Visual representation of the “8-cube” dataset across 8 genotypes, 8 tissues, and 8 replicates. **b**, Experimental design of sample barcoding plate. Nuclei derived from one tissue serves as the main tissue on the plate, where each sample has its own unique sample well and corresponding barcode. Nuclei from a different tissue are multiplexed in the remaining third of the plate, where two samples from distinct genotypes for the same sex and replicate are loaded into one well. **c**, Tissue loading pattern for all 8 sample barcoding plates. **d**, Pseudobulk PCA of 516 total samples colored by tissue and points indicating females (“x”) and males (circle). **e**, Adjusted p-values from ANOVA test of significance between PCs and metadata (top); scores for the first 9 PCs across samples (bottom). **f**, Normalized expression of 20 protein-coding genotype-specific genes across pseudobulk samples grouped by tissue, genotype, and sex. **g**, Number of up- and down-regulated differentially expressed genes (adj. p-value < 0.01, |LFC|>1) between each genotype and B6. **h**, Percentage of nuclei falling into genotype-specific clusters across tissues.

Nuclei were extracted from each tissue in the 8-cube dataset, with a few additional replicates increasing the total number to 516 samples from 73 mice (Methods). We performed combinatorial split-pool barcoding experiments to build single-nucleus RNA-seq libraries ^51,52^. In this approach, fixed nuclei are first distributed into a 96-well plate, with each well containing unique barcodes that are appended to RNA during reverse transcription to record the sample identity. Each RNA-seq library contains a limited number of cells (between 12,000 and 65,000) that we refer to as subpools, and each experimental plate produces multiple subpools. To detect experimental batch effects and maximize sample sequencing, we multiplexed samples in one-third of each experiment (Fig. 1b). In the first eight columns of the plate, each well contains a unique sample from a single tissue. The remaining four columns contain samples from a second tissue, with each well multiplexing two samples from distinct genotypes. However, with this approach, we observed that multiplexing tissues on the same barcoding plate introduced detectable RNA contamination between neighboring tissues. For example, differential expression analysis between our B6 data and a supplemental dataset from a mouse model of Alzheimer’s disease with a B6 background, *Trem2*^R47H NSS^ homozygotes (“Trem2”), revealed substantial batch effects linked to the inclusion of all 8 tissues plus PBMCs on the sample loading plate (Supp. Fig. 2a-b). This suggested that some degree of batch effect would be present throughout the 8-cube dataset. To mitigate this, we applied CellBender ^53^, a software tool designed to remove technical artifacts caused by ambient RNA contamination, PCR chimeras, and other sources of noise (Methods, Supp. Fig. 2c). CellBender was applied to each subpool library in our 8-cube and Trem2 datasets, substantially reducing cross contamination between tissues (Methods, Supp. Table 2). This removal of tissue-specific ambient signals improved the accuracy of downstream analyses, enabling more confident detection of true biological differences across genotypes (Supp. Fig. 2d-e).

**Figure 2.**
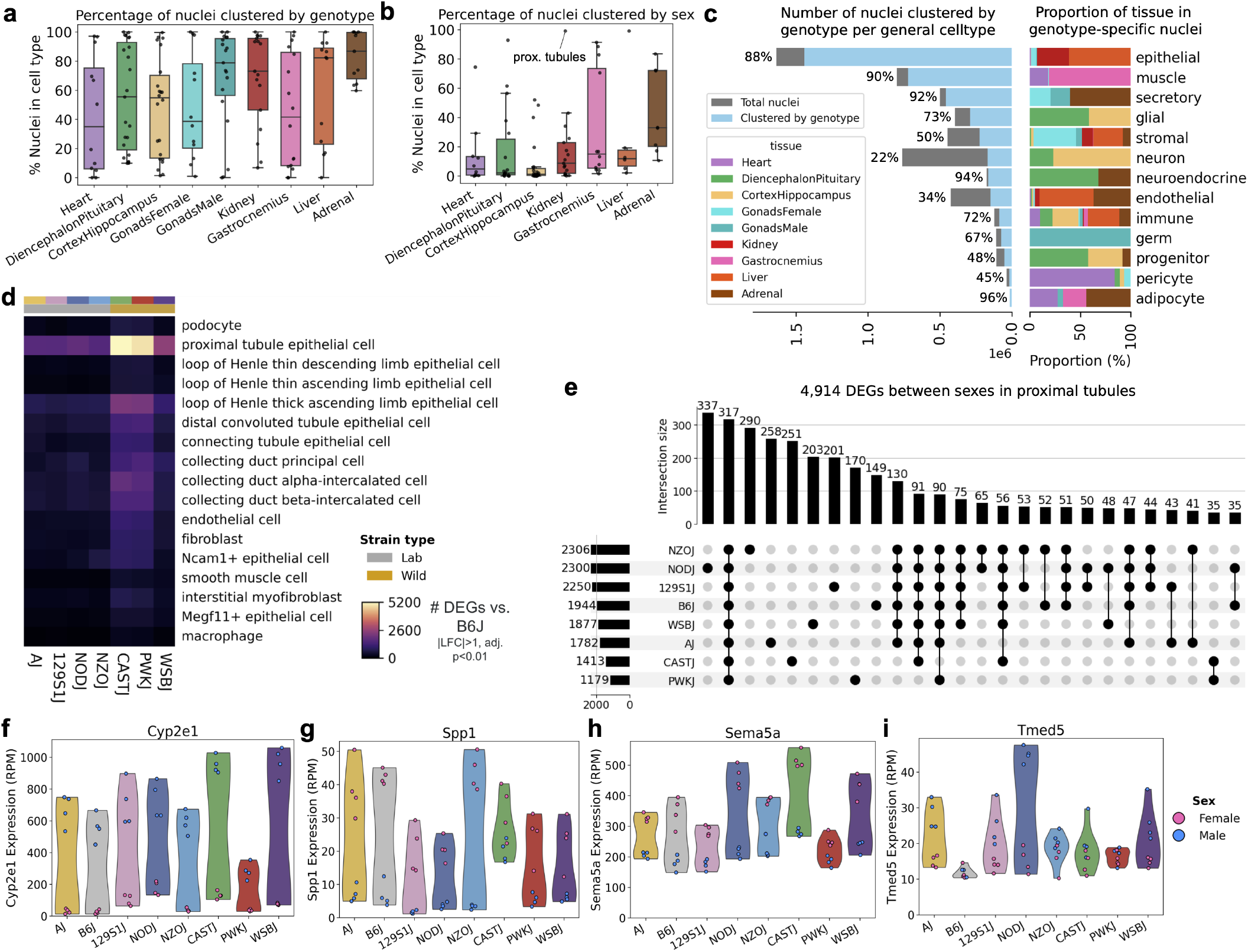
Genotype- and sex-driven clustering across cell types and general cell type lineages. **a**, Distribution of the percentage of nuclei per cell type assigned to genotype-specific subclusters, grouped by tissue of origin. **b**, Distribution of the percentage of nuclei per cell type assigned to sex-specific subclusters, excluding gonadal cell types. **c**, Proportions and counts of nuclei clustering by genotype across 13 general cell lineages, showing tissue distribution of nuclei in genotype-specific clusters. **d**, Number of DEGs in each cell type in kidney (both up- and down-regulated) in each genotype compared to B6 (adj. p-value < 0.01, |LFC|>1). **e**, Overlap of DEGs between sexes within each genotype in kidney proximal tubule epithelial cells (adj. p-value < 0.01, |LFC|>1). f-i, Pseudobulk expression of DEGs in kidney proximal tubule epithelial cells showing genotype-specific variation by sex: **f**, Male-upregulated *Cyp2e1*. **g**, Female-upregulated *Spp1*. **h**, Female-upregulated *Sema5a*. **i**, Male-upregulated *Tmed5*.

To design the sample multiplexing scheme, we calculated pairwise Hamming distance between strains based on 1,537,904 SNPs in 3’ UTRs and exons, followed by maximal weight matching to pair the most genetically distinct strains. This resulted in the following pairings: CAST with 129S1, NOD with B6, PWK with AJ, and WSB with NZO (Fig. 1b). Multiplexed wells required genetic demultiplexing to correctly assign nuclei to their strain of origin (Methods). The multiplexing scheme was repeated for all eight tissues (Fig. 1c). Each plate corresponds to an experiment with several library subpools and an expected yield of 1 million nominal nuclei conducted using Parse Biosciences Evercode Mega kits ^52^ (Methods). Thus, for each tissue and each mouse replicate, samples were sequenced twice, once as an individual sample and once multiplexed with another genotype (Fig. 1c). We sequenced approximately 1 million nuclei per tissue with 20 billion reads for a nominal depth of 20,000 reads per nucleus and aligned using mm39 with GENCODE vM32 annotations (Methods). The final dataset spans 8 million nominal nuclei sequenced with 165 billion short reads. After quality control filtering (Methods), we retained 5.2 million nuclei, 620,258 from B6 alone. Across all genotypes, we recovered 794,625 nuclei from muscle, 633,651 from liver, 649,423 from kidneys, 442,018 from ovary and oviduct, 255,862 from testes and epididymis, 638,843 from cortex and hippocampus, 805,861 from diencephalon and pituitary gland, 509,714 from adrenal glands, and 458,167 from heart.

### Tissue-independent gene expression associated with genetic variation

A pseudobulk principal component analysis (PCA) across all data at the sample level revealed high replicate concordance and expected clustering patterns between tissues (Fig. 1d). PC1 (31.59% of the variance) separated ectoderm-derived brain regions in positive PC1 from endoderm-derived kidney and liver in negative PC1, while PC2 (18.41% of the variance) separated mesoderm-derived heart and muscle tissues. Female gonads and adrenal glands clustered together in PCs 1 and 2, but were separated in later PCs (Fig. 1e), highlighting the shared function of these organs as key components of the endocrine system. Analysis of the top 9 PCs encompassing 90% of the total variance in the dataset revealed significant associations between tissue and PCs 1-8, and between sex and PCs 3, 4, and 7. Interestingly, PC9 (1.04% of the variance) was strongly associated with genotype rather than tissue or sex by separating the *domesticus* strains from non-*domesticus* (PWK and CAST) strains. PCA of samples by each tissue separately revealed genotype-driven patterns, where non-*domesticus* samples clustered apart from the other six genotypes in either PC1 or PC2 accounting for 14-26% of the total variance (Supp. Fig. 3a-i). Although cross-tissue expression patterns were primarily driven by tissue and sex, deeper analysis at the level of individual tissues can reveal substantial genotype effects, reinforcing the importance of resolution in identifying genotype-specific variation.

**Figure 3.**
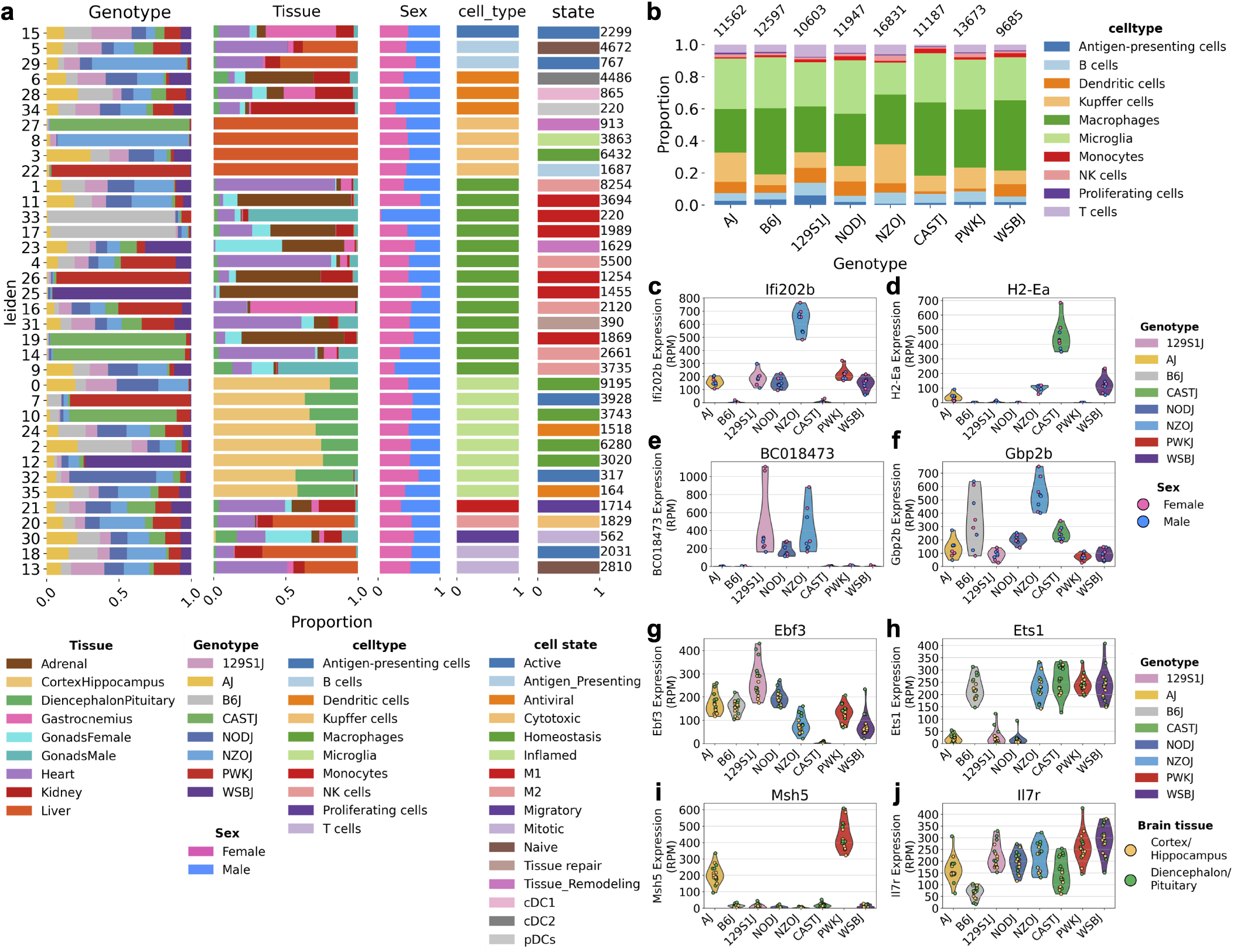
Pan-tissue analysis of immune cell composition and gene expression differences across genotypes. **a**, Proportion bar plots showing the breakdown of genotype, tissue, sex, cell type, and cell state across all 36 immune cell clusters (98,085 nuclei). Each row corresponds to an individual cluster and each column represents a different categorical variable, with cluster number indicated on the left and total nuclei per cluster on the right. **b**, Stacked bar plots depicting the proportional composition of immune cell types across each genotype. c-f, Pseudobulk expression of genes in liver Kupffer cells: **c**, *Ifi202b*, **d**, *H2-Ea*, **e**, *BC018473*, **f**, *Gbp2b*. g-j, Pseudobulk expression of genes in microglia from both brain regions: **g**, *Ebf3*, **h**, *Ets1*, **i**, *Msh5*, **j**, *Il7r*.

We further characterized genotype-driven variation in the dataset by variance decomposition (Methods), which identified 82 genes that varied more by genotype than by tissue. We found 26 genes were consistently upregulated in CAST, 24 in PWK, 18 in WSB, 5 in B6, 3 in 129S1, and 2 each in AJ, NZO, and NOD (Supp. Table 3). Of the 82 genes, half are pseudogenes, 20 are protein-coding (Fig. 1f), 16 are lncRNAs, 2 are transcribed enhancer RNAs, 1 is a ribozyme (*Rpph1* in B6), and 1 is a microRNA (*Mir1291* in CAST). Interestingly, several of the 20 protein-coding genes encode chemoreceptive G protein-coupled receptors, for example olfactory receptor *Or1p1c* in NOD, taste receptor *Tas1r1* in CAST, and vomeronasal receptors *Vmn1r196* and *Vmn1r195* in PWK as well as *Vmn2r66* in WSB. This suggests that sensory or environmental interactions may be regulated differently across strains. Chemoreceptive receptors, such as those involved in detecting odors and tastes, may contribute to strain-specific behaviors or ecological adaptations, highlighting potential differences in sensory biology and environmental responses between the strains and subspecies.

Beyond sensory genes, several other strain-specific genes suggest metabolism-related adaptations. A CAST-specific gene, *Kcnmb3*, encodes the auxiliary beta subunit of calcium-activated potassium channels, which are known for regulating cell excitability and neurotransmitter release ^54^. In WSB, the cytokine gene *Il36g* was highly expressed compared to other genotypes. Interestingly, prior work showed that WSB mice have lower circulating levels of several inflammatory cytokines, including IL-5, IL-6, and IL-10, contributing to their resistance to diet-induced obesity ^55^. This suggests that WSB may rely on distinct cytokine programs compared to other strains, highlighting how strain-specific immune responses can shape metabolic phenotypes. In diabetes-prone NZO mice, we also observed body-wide over-expression of *Gjb4*, which encodes a gap junction protein. Overexpression of *Gjb4* in NZO pancreatic islets has been shown to inhibit islet proliferation and reduce insulin secretion, contributing to islet dysfunction and type 2 diabetes ^56^. In 129S1, the gene *Klk1b22* was specifically expressed. Increased expression of this gene has been detected in circulating 129S1 T cells, which has been linked to their improved glucose tolerance compared to B6^57^. Finally, WSB showed striking overexpression of *Cwc22*, which encodes a spliceosome-associated protein ^58^. This overexpression directly reflects a copy number expansion at the R2d2 locus, located approximately 6 Mb downstream from its paralog R2d1^59,60^. *Cwc22* is the only protein-coding gene within this region, and amongst the founder strains its expression is uniquely amplified in WSB ^60^. Together, these findings suggest that strain-specific gene expression may contribute to differences in immune regulation, metabolism, and other physiological traits.

### Genotypic impact on gene expression varies across tissues

We performed differential expression analysis between every pairwise comparison in each tissue at the pseudobulk level in order to identify additional differences that could be tissue-specific. The highest number of differentially expressed genes (DEGs) was consistently observed in pairwise comparisons involving non-*domesticus* strains CAST and PWK (Supp. Fig. 4a). To align with the broader community’s emphasis on B6 as the reference strain for mouse models and the reference genome, we focused our primary analysis on differences between B6 and each of the wild-derived strains (Fig. 1g). Male gonads had the highest number of DEGs overall, likely reflecting the broader transcriptional landscape of germ cells. Among the 3,303 DEGs identified exclusively in male gonads in B6 comparisons (Supp. Fig. 4b), 32% were protein-coding, 33% were lncRNAs, and 28% were pseudogenes. Among the male gonads-specific proteincoding DEGs, 26% were olfactory receptors and 5.2% were vomeronasal receptors, potentially reflecting the specialized roles of these genes in processes such as sperm chemotaxis ^61^. Most DEGs were either shared across multiple tissues or restricted to a specific tissue (Supp. Fig. 4b). Only 4.1% of DEGs were found in every tissue, including 66 of the 82 genes identified in the variance decomposition analysis from the previous section. Male gonads had the highest percentage of tissue-specific DEGs, followed by liver (Supp. Fig. 4c). Interestingly, 13% of genes upregulated in WSB are specific to adrenal tissue, compared to 5.3% in other genotypes (Supp. Fig. 4c). This trend was also reflected in the overall number of DEGs, with WSB having 1.9 times more adrenal DEGs than other *domesticus* strains (Fig. 1g). GO biological process and Reactome terms associated with 801 upregulated genes in WSB compared to B6 specifically in adrenal included neuronal system, axonogenesis, regulation of monoatomic cation transmembrane transport, and norepinephrine biosynthetic process (p < 0.05, Supp. Fig. 4d). NZO showed the most significant GO term enrichment among 405 adrenal-specific upregulated genes, with terms including cholesterol biosynthesis, sterol biosynthesis, and secondary alcohol biosynthetic process (Supp. Fig. 4e). Notably, genes coding for apolipoproteins such as apolipoprotein B (*Apob*) and apolipoprotein A2 (*Apoa2*) were upregulated in NZO adrenals. Variants in Apob have been linked to obesity, high cholesterol, and increased risk of coronary heart disease in men ^62^, and serum ApoB levels correlate with inflammatory markers in obese women ^63^. Additionally, elevated serum ApoB has been associated with increased risks of metabolic syndrome, hypertension, and diabetes, independent of central obesity and inflammation markers ^64^. These findings suggest that upregulation of *Apob* in NZO adrenal glands may contribute to or result from the strain’s obesity phenotype, potentially reflecting physiological responses to obesity rather than direct effects of genetic variation.

**Figure 4.**
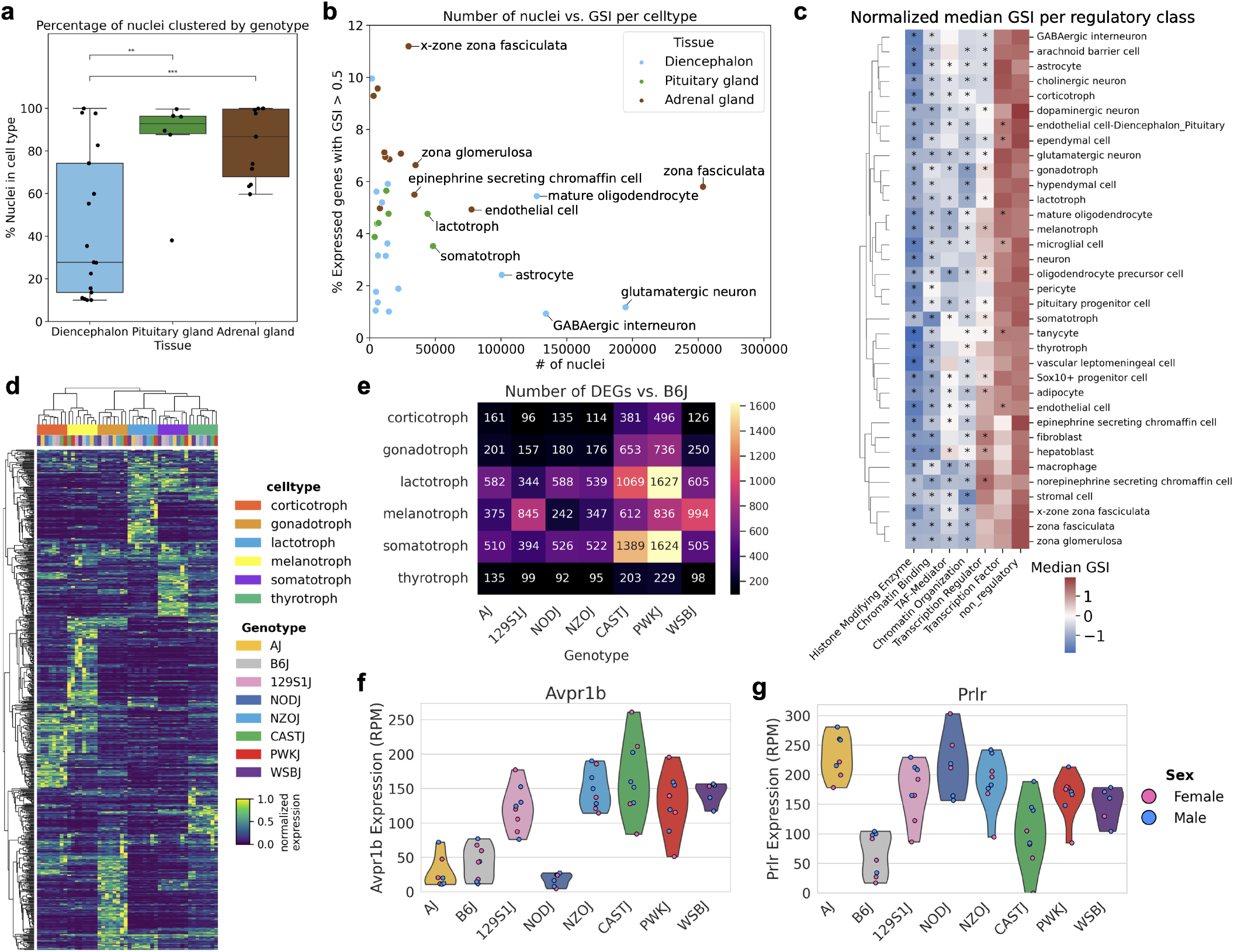
Genotype-specific gene expression in tissues of the HPA axis. **a**, Proportion of nuclei clustering into genotype-specific clusters in the diencephalon, pituitary gland, and adrenal gland, with asterisks indicating significance (p < 0.05, t-test). **b**, Comparison of the total number of nuclei and the percentage of genes with a genotype specificity index (GSI) > 0.5 across cell types. **c**, Row-normalized median GSI values centered around 0 for different gene regulatory classes, with asterisks indicating significant differences (p < 0.05, Kolmogorov–Smirnov test) relative to non-regulatory genes. **d**, Heatmap of 720 genes with GSI > 0.5 in at least one pituitary gland cell type. **e**, Total number of differentially expressed genes compared to B6 in six pituitary gland cell types. f-g, Pseudobulk expression of genes in corticotroph nuclei: **f**, *Avpr1b* and **g**, *Prlr*.

When annotating cell types, we observed that some cell types formed a single unified cluster, while others consisted of multiple subclusters, each corresponding to a different genotype and/or sex. To quantify the impact of genotype on single-nucleus clustering, each cell type within each tissue was systematically subclustered using the Leiden algorithm at default resolution (Methods). To minimize pan-tissue effects, global genotype-specific genes identified in the previous variance decomposition analysis were excluded from the set of highly variable genes used for clustering. We then compared the genotype distribution within each subcluster to the overall distribution of genotypes in the cell type, identifying genotype-specific subclusters (chi-squared p < 10^−10^, Methods). Finally, we calculated the proportion of nuclei in each genotype and tissue that fell into genotype-specific subclusters (Fig. 1h). Across strains, the majority of nuclei in gonads, kidney, muscle, liver, and adrenal gland were grouped into genotype-specific subclusters. More than 50% of CAST and PWK nuclei cluster separately from the *domesticus* strains in all tissues except for the heart, a trend that mirrors the differential expression results, where the heart is least impacted by genotype. However, this clustering-based approach de-emphasizes factors such as the total number of genes expressed and cell type heterogeneity, which particularly affected DEG counts in brain tissues. Instead, it captures the extent to which genotype influences the selection of highly variable genes used for clustering. While DEG numbers correlate strongly with cell type size due to UMI coverage (R^2^ = 0.89, average across tissues, Supp. Fig. 5a), this clustering-based analysis highlights the magnitude of genotype-driven expression differences in the most variable genes and remains independent of cell size. The conservation of gene expression in heart and brain tissues suggests that core physiological processes in these organs are less susceptible to genetic variation. In contrast, tissues such as the liver and adrenal glands that are highly responsive to environmental cues, hormones, and metabolic demands exhibit greater genotype-driven expression differences. Notably, WSB nuclei clustered at levels similar to other *domesticus* strains, except in the diencephalon/pituitary, male gonads, and liver, where its wild-derived background may have a greater impact. In contrast, the non-*domesticus* strains PWK and CAST exhibited a 1.5-fold increase in genotype-specific clustering compared to *domesticus* strains, particularly in brain regions and female gonads (Fig. 1h). This analysis demonstrates how genotype differentially influences single-cell clustering across tissues, with stronger effects observed in metabolically active and hormone-responsive tissues.

**Figure 5.**
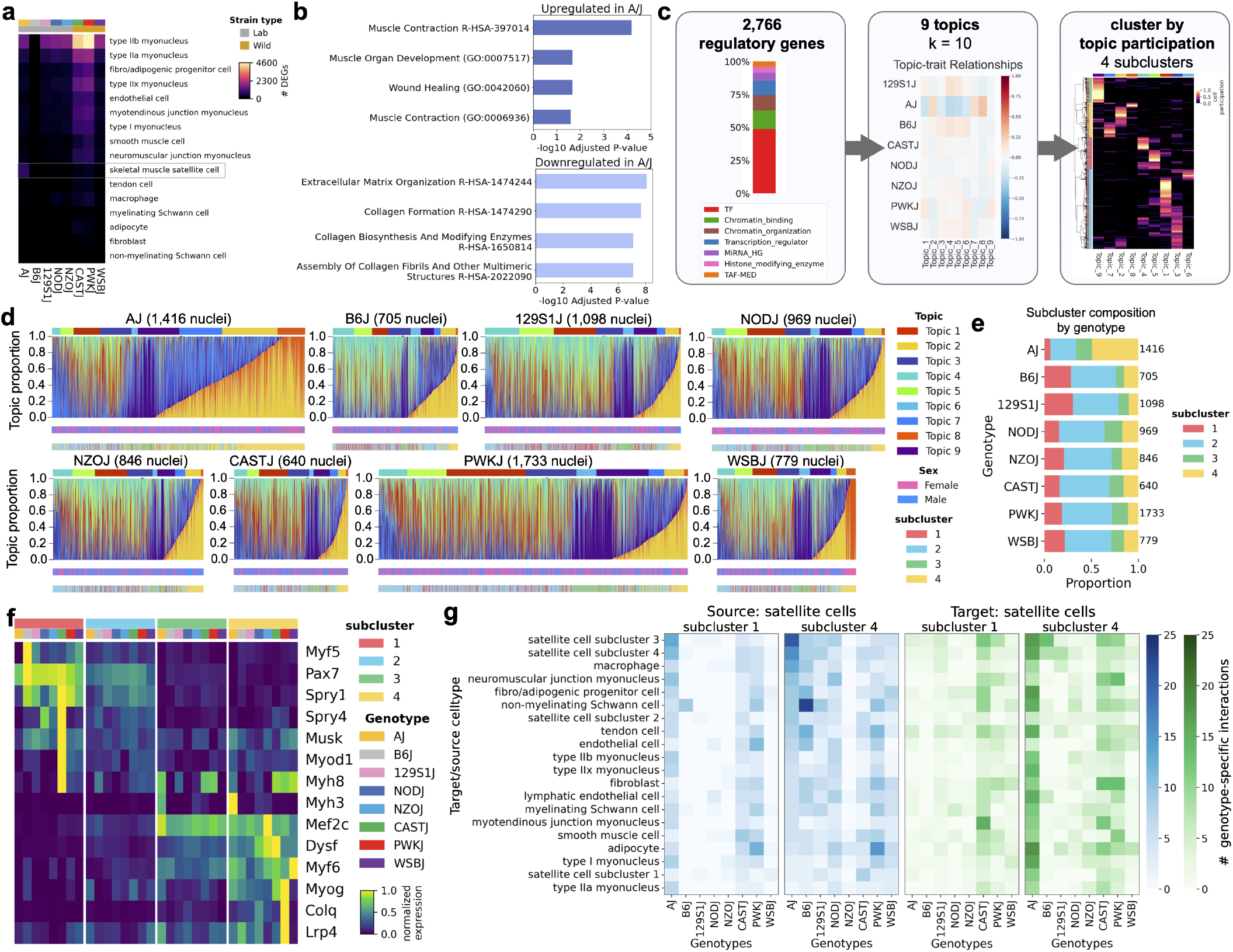
Satellite cells are over-activated in AJ. **a**, Number of differentially expressed genes between each genotype and B6 (adj. p-value < 0.01 and |LFC| > 1) in skeletal muscle cell types. **b**, GO biological process and Reactome terms for genes upregulated and downregulated in AJ compared to B6 (adj. p-value < 0.05). **c**, Overview of topics modeling analysis on 8,186 satellite cells. **d**, Structure plots of topic proportions per nucleus grouped by genotype. Each column is a stacked bar plot showing the proportion of participation across the 9 topics for each nucleus across 8,186 total nuclei in satellite cells. **e**, Distribution of nuclei across subclusters determined by clustering cells by participation across topics. **f**, Heatmap of pseudobulk expression in satellite cells grouped by subcluster and genotype. **g**, Number of unique interactions between cell types and subtypes, including satellite cell subclusters, where subclusters 1 and 4 are the source (blue) and when subclusters 1 and 4 are the target (green).

### Cell-type-specific effects of genotype and sex

Analyzing the percentage of nuclei clustering by genotype revealed a broad range of genotype-driven effects at the cell type level (Fig. 2a, Supp. Fig. 5b-j). In every tissue, particularly adrenal, several cell types exhibited significant genotype-specific clustering. For example, a majority of nuclei (>50%) clustered by genotype in 6 of 12 heart cell types, 12 of 23 diencephalon/pituitary cell types, 13 of 22 cortex/hippocampus cell types, 5 of 12 female gonad cell types, 15 of 19 male gonad cell types, 7 of 16 muscle cell types, 7 of 12 liver cell types, and 12 of 17 kidney cell types (Fig. 2a). Notably, all 11 adrenal cell types showed genotype-driven clustering. This suggests that while not all cell types within a tissue are affected, genotype influences at least some cell types across all tissues. A similar analysis was conducted for sex-specific clustering, where nuclei were subclustered within each cell type and genotype, and sex-specific clusters were identified using a chi-squared test (p < 10^−10^, Methods). This approach captures the magnitude of sex-specific clustering independently of genotype. Here, significant sex-driven clustering, where more than 50% of nuclei in a given cell type fall into a sex-specific cluster, was observed in ventricular cardiac myocytes, all myonuclear subtypes in skeletal muscle, hepatocytes, kidney proximal tubule epithelial cells, and three adrenal cell types (zona fasciculata, zona glomerulosa, and epinephrine-secreting chromaffin cells) (Fig. 2b). In cortex/hippocampus, most cell types exhibited minimal sex-specific clustering (<20% of nuclei in the cell type), except for choroid plexus epithelial cells (52%), dentate gyrus granule cells (40%), and oligodendrocyte precursor cells (48%). Consistently, oligodendrocyte precursor cells (OPCs) also showed sex-specific clustering (38%) in the diencephalon/pituitary. Prior studies have highlighted sex-specific differences in cellular characteristics and gene expression in OPCs, along with differential regulation of proliferation by sex hormones ^65,66^. Interestingly, all pituitary-specific cell types clustered by sex, with the majority of gonadotrophs (87%) and lactotrophs (93%) falling into sex-specific clusters (Fig. 2b). This analysis demonstrates that while genotype strongly influences cell type clustering across all tissues, its effects are concentrated in specific cell types within each tissue, whereas sex-specific effects are more restricted to an even smaller subset of cell types.

Examining genotype-driven clustering within broader cell lineages revealed distinct trends (Fig. 2c, Supp. Fig. 5k). The majority of nuclei in the most abundant cell type in the dataset, epithelial cells (1.6 million nuclei) clustered by genotype and sex, predominantly in kidney proximal tubule cells and liver hepatocytes. Muscle cells (797k nuclei) exhibited significant clustering by both genotype and sex in heart and gastrocnemius. In contrast, neurons (761k nuclei) were the least impacted by genotype, with more nuclei clustering by genotype in cortex/hippocampus compared to diencephalon, particularly in dentate gyrus, CA1, and CA3 hippocampal neurons (Supp. Fig. 5b). A key consideration of this analysis is that cell type complexity may obscure genotype and sex effects at this resolution. Higher-resolution subclustering could potentially reveal additional genotype- and sex-specific patterns, particularly in cell types such as neurons where inter-cell type heterogeneity plays a larger role in clustering than genotype or sex differences. In contrast, cell types with more homogeneous populations, such as hepatocytes, proximal tubule epithelial cells, and skeletal and cardiac muscle more clearly exhibited genotype and sex effects. Because these populations lack finer subtypes beyond our annotations, the remaining variation is largely attributable to genotype and sex differences. This analysis does not measure the strength of these effects but instead demonstrates their presence, enabling comparisons across cell types independent of cell type abundance. Overall, epithelial, muscle, secretory, neuroendocrine, and adipocyte cell types are the most affected by genotype and sex. In contrast, glial and immune cells exhibit more genotype-specific clustering than sex-specific clustering, highlighting differential susceptibility to genetic background versus sex differences across cell types.

### Renal tubules show pronounced genotypic and sex differences

In the kidney, all renal tubule epithelial cell types exhibited strong genotype-specific clustering. As the largest cell type at 303k nuclei, proximal tubule epithelial cells were also the only renal cell type to show significant sex-specific clustering (Fig. 2b). The next largest epithelial population was the loop of Henle thick ascending limb epithelial cells which was similar in size to endothelial cells (77,123 nuclei compared to 64,785 nuclei), yet endothelial cells exhibited much less genotype-specific clustering (Supp. Fig. 5f). This pattern was consistent across tissues, where endothelial cells were among the least affected by genotype-specific clustering, except in the liver and adrenal gland (Fig. 2c, Supp. Fig. 5b-j). To further investigate these differences, we identified DEGs between each genotype and B6 within each kidney cell type (Fig. 2d, Methods). As expected based on cell numbers, proximal tubule epithelial cells had the highest number of DEGs. Similar to broader tissue-level analyses, most DEGs were detected in comparisons involving CAST and PWK across all kidney cell types. Notably, collecting duct alpha and beta principal cells have a high number of DEGs relative to their size. Unlike loop of Henle cell types, genotype-specific clustering was observed in every strain in collecting duct principal cells, not just primarily in CAST and PWK (Fig. 2d, Supp. Fig. 5f). This may reflect genetic variation in hormone responsiveness, as these cells regulate sodium and water balance in response to vasopressin and aldosterone ^67^. More broadly, genetic variation in renal epithelial cells may underlie strain-specific differences in kidney function, influencing solute transport, fluid homeostasis, and drug metabolism.

Proximal tubule epithelial cells, identified in previous mouse and human single-cell studies as uniquely impacted by sex ^68,69^, were the only kidney cell type to show strong sexspecific clustering in this analysis (Fig. 2b). To explore this further, we compared female to male proximal tubule cells within each genotype and identified 4,914 unique DEGs upregulated in either females or males across genotypes (Fig. 2e). While 317 genes were differentially expressed between sexes across all genotypes, 1,859 were unique to a specific genotype. Of 60 previously identified sex-specific genes ^68^, 53 were differentially expressed in at least one strain, including female-upregulated *Slc7a12* and male-upregulated *Cyp7b1* and *Cyp2e1* (Fig. 2f), which were detected in all eight genotypes. Interestingly, sexual dimorphism of gene expression was absent in certain genotypes. For example, *Spp1* (osteopontin), linked to kidney fibrosis, inflammation, and disease, is regulated by sex hormones ^70–73^, and was differentially upregulated in females in every genotype except CAST (Fig. 2g). It is important to acknowledge that some genes may appear unique to certain genotypes not because they lack differential expression in others, but because their expression differences do not exceed our log fold change threshold. Even among genes detected as differentially expressed in multiple genotypes, variations in expression magnitude were notable. For example, *Cyp2e1* was differentially expressed in all genotypes, yet its male-upregulated expression was significantly reduced in PWK (Fig. 2f). NOD had the highest number of unique DEGs, with 337 genes. For example, *Sema5a* was upregulated only in NOD females, but also showed similar sexually dimorphic expression in most strains except PWK (Fig. 2h). Similarly, *Tmed5* was significantly upregulated in NOD males and also exhibited differences in expression between sexes in AJ, 129S1, and WSB (Fig. 2g). *Sema5a* has been previously identified as downregulated in glomerular tissues of human diabetic kidney disease patients ^74^, a condition where men with both type 1 and type 2 diabetes are at higher risk than women ^75^. *Tmed5*, a transmembrane trafficking protein, has been linked to hepatocellular carcinoma, where its overexpression promotes tumor progression ^74^. Together, these findings highlight strain-dependent variation in sex-biased gene expression in proximal tubules, reinforcing the interplay between genetic background, sex, hormone regulation, and disease susceptibility in the kidney.

### Pan-tissue analysis of immune cells reveals genotype--driven trends

The only cell types shared across all tissues were endothelial cells and immune cells (Fig. 2c). Given that the majority of immune cells exhibited genotype-specific clustering, along with prior studies in CC founder strains showing differences in immune cell composition in blood, such as varying proportions of T cells, NK cells, and dendritic cells ^76^, we conducted a pan-tissue immune-focused analysis. We aggregated all immune cells (98,085 nuclei) across tissues, which was composed of 35% resident macrophages (34,770 nuclei), 13% liver Kupffer cells (specialized resident macrophages, 12,895 nuclei), 29% microglia from both brain regions (28,165 nuclei), and lymphocytes such as B cells and T cells (10,280 total nuclei). Clustering all immune cells together across tissues allowed for finer annotation of immune subtypes that were indistinguishable within a single tissue. We identified 36 clusters (Fig. 3a) spanning all eight tissues, enabling the annotation of 10 distinct immune cell types, including smaller populations such as antigen-presenting cells (2,299 nuclei), dendritic cells (5,571 nuclei), monocytes (1,714 nuclei), NK cells (1,829 nuclei), and proliferating immune cells (562 nuclei). Among these 36 clusters, half exhibited significant genotype-specific clustering (chi-squared p-value < 10^−10^). These included six microglia clusters, where CAST, PWK, WSB, and NOD form distinct clusters, three Kupffer cell clusters specific to CAST, PWK, and NZO, seven macrophage clusters specific to CAST, PWK, and WSB, and one B cell cluster specific to NZO. Additionally, a small cluster (cluster 33, 220 nuclei) was composed primarily of B6 male macrophages. However, with 67% of the nuclei derived from a single replicate, its biological relevance is uncertain. Sex was evenly distributed across the remaining clusters, consistent with findings from per-tissue clustering analyses (Supp. Fig. 5k).

We further annotated functional states across immune cell types. In Kupffer cells (KCs), we identified cluster 8 as an NZO-specific inflamed state. This cluster comprises 3,863 nuclei, representing 86% of total NZO KCs. As the largest population of tissue-resident macrophages in the body, KCs are derived from embryonic yolk sac progenitors and complete differentiation in the fetal liver where they establish a self-renewing population ^77^. Studies suggest the existence of two phenotypically and functionally distinct populations of KCs: a major CD206-low and ESAM-negative population (KC1) and a minor CD206-high ESAM-positive population (KC2). Both subsets express core markers such as *Clec4f, Vsig4, Csf1r*, and *Adgre1* (Supp. Fig. 6a) ^78^. KC1 cells are primarily associated with immune functions, while KC2 cells exhibit higher expression of endothelial markers and genes linked to metabolic pathways, including lipid metabolism ^78,79^. In our dataset, all KC clusters except cluster 27, which is CAST-specific, expressed *Mrc1* but lacked *Esam* expression, suggesting that they do not match the described KC2 profile (Supp. Fig. 6a). Cluster marker gene analysis followed by GO and Reactome term enrichment revealed that genes defining the NZO-specific KC cluster are enriched for the innate immune system, regulation of phagocytosis, and regulation of TNF cytokine production (Supp. Fig. 6b). This cluster exhibits upregulation of genes such as *Tlr8, Irak3, Mefv, Il17ra*, and *Pla2g7* (Supp. Fig. 6c), supporting an activated inflammatory state. *Irak3* is a modulator of inflammation ^80^, while IL-17 signalling in KCs has been implicated in liver fibrosis ^81^, suggesting that the inflammatory response in NZO KCs may contribute to liver pathology. Given these findings, this cluster likely represents an activated inflammatory KC population. Rather than purely reflecting genetic differences, this inflammatory response may be driven by the severe fatty infiltration observed in NZO livers, which were twice the size of those from other genotypes (Supp. Fig. 1i).

We also observed differences in immune cell proportions that align with known strain-specific trends (Fig. 3b) ^76^. For example, CAST had 4.8-fold fewer T cells compared to other genotypes, while NZO had 2.5 times more NK cells. Additionally, 129S1, WSB, and NOD had 2.2-, 1.8-, and 2.2-fold higher proportions of dendritic cells than the other five genotypes, respectively (Fig. 3b). NZO showed a 2.3-fold expansion of Kupffer cells compared to other genotypes. Examining differentially expressed genes in KC clusters revealed several genotype-specific genes of interest. Notably, *Ifi202b* (IFI16), an interferon-activated gene linked to obesity and adipogenesis in mice and humans ^82^, was identified as a marker gene for KC cluster 8 and is highly expressed specifically in NZO (Fig. 3c). Its overexpression promotes adipogenesis and fat storage, contributing to obesity-associated insulin resistance ^82^. *H2-Ea*, a class II major histocompatibility complex (MHC) gene, was highly expressed in CAST (Fig. 3d). Given that KCs are known to express both MHC class I and II molecules ^83^, *H2-Ea* may play a key role in liver immunity and antigen presentation to T cells. We detected a long noncoding RNA (lncRNA) *BC018473* which was highly expressed in 129S1, NOD, and NZO, but nearly absent in other genotypes including B6 (Fig. 3e). This gene is uncharacterized but has been identified as differentially expressed in CD4 T-cells between mice from B6 and NOD backgrounds, with higher expression in NOD ^84^. Another highly specific gene, *Gbp2b*, codes for a guanylate-binding protein involved in innate immunity ^85^ and is uniquely expressed in NZO, with variable expression in B6 (Fig. 3f). *Gbp2b* plays a role in interferon-induced responses to pathogens and has been associated with inflammation ^86,87^. These findings illustrate the substantial genotype-driven variability in KCs, suggesting strain-specific differences in immune function, inflammation, and metabolic adaptation. In particular, NZO KCs form a distinct cluster characterized by interferon gene activation, which may reflect an inflammatory response driven by the metabolic consequences of obesity rather than purely genetic variation.

Finally, we examined differential gene expression in microglia, where we identified the largest genotype-specific clusters. Microglia had 2.5-fold as many nuclei falling into genotype-specific clusters compared to macrophages. Among the differentially expressed genes, we identified key immune-related transcription factors, including *Ebf3* and *Ets1. Ebf3*, a transcription factor involved in neuronal and immune regulation, varied in expression across the eight genotypes and was particularly low in CAST (Fig. 3g). *Ebf3* and its antisense lncRNA, *Ebf3-AS*, have been linked to neurodevelopmental disorders and neuroinflammation, particularly in Alzheimer’s disease models, where they influence microglial function and apoptosis ^88^. *Ets1* was highly expressed in microglia from five genotypes, including B6, but showed low expression in AJ, 129S1, and NOD (Fig. 3h). This transcription factor regulates differentiation, survival, and cytokine expression in immune cells, including microglia ^89,90^. We also detected *Msh5*, a gene typically associated with meiosis ^91,92^, showing differential expression in microglia (Fig. 3i). Although its specific function in microglia is unclear, *Msh5* has been linked to immune deficiencies and ovarian disorders ^93,94^, suggesting a potential broader function in immune regulation. Additionally, *Il7r*, a key receptor for IL-7 signaling, was more lowly expressed in B6 than the other strains (Fig. 3j). IL-7 is primarily known for its role in T and B cell homeostasis but has also been detected in microglia under certain conditions ^95^. Notably, IL-12 has been shown to induce IL-7 expression in both mouse and human microglia, suggesting a potential role in neuroimmune interactions ^95^. The expression patterns of these genes were consistent across both brain regions, reinforcing a genotype-driven effect on microglial gene regulation. Overall, these findings reveal extensive genotype-driven variation in immune cell composition, inflammatory states, and gene expression, underscoring the role of genetic background on immune regulation and potential disease susceptibility across strains.

### Transcriptional variation in distinct cell types of the HPA axis

The hypothalamus-pituitary-adrenal (HPA) axis is a major endocrine system involved in stress response and regulation of metabolic, immune, and cognitive processes through glucocorticoid signaling ^96^. In response to stressful stimuli, sensory input reaches the paraventricular nucleus (PVN) of the hypothalamus, leading to activation of corticotropin-releasing hormone (CRH) neurons and CRH release ^97^. CRH travels to the pituitary, where it stimulates corticotrophs to release adrenocorticotropic hormone (ACTH). ACTH, in turn, induces release of glucocorticoids such as cortisol (corticosterone in mice) from the zona fasciculata of the adrenal glands ^96^. The HPA axis is highly plastic and responsive to environmental cues and prior experience ^98,99^. The hypothalamus regulates many aspects of bodily homeostasis, including thermoregulation, hunger, circadian rhythms, and reproductive functions through coordinated hormonal signaling with the pituitary gland ^100^. Beyond its neuroendocrine functions, the hypothalamus also integrates signals from various brain regions, influencing behaviors related to homeostasis, behavior, and stress ^100,101^. While our dissections focused on recovering primarily hypothalamic tissue, we also captured neighboring thalamic regions, which are characterized by high *Tcf7l2* expression ^102^ (Supp. Fig. 7a), as well as the pituitary gland. Of the 23 cell types identified in the diencephalon and pituitary gland (805,861 total nuclei), six were neuroendocrine cell types from the pituitary, representing 15% of nuclei (121,624 total nuclei). The pituitary gland is anatomically divided into three main sections, where the anterior lobe is the largest ^103^. This region contains five primary hormone-secreting cell types: somatotrophs, which produce growth hormone; lactotrophs, which secrete prolactin; thyrotrophs, responsible for thyrotropin production; corticotrophs, which release ACTH; and gonadotrophs, which secrete gonadotropins, including follicle-stimulating hormone and luteinizing hormone. The intermediate lobe contains the sixth cell type, melanotrophs, which produce melanocyte-stimulating hormone ^103^. Analysis of the six pituitary cell types separately from those in the diencephalon revealed clustering by genotype was significantly stronger in pituitary. By contrast, the adrenal gland exhibited a broader range of genotype-specific clustering across its cell types, but still showed significantly more than the diencephalon (Fig. 4a). These findings suggest that genotype more strongly influences transcriptional variation in endocrine tissues, such as the pituitary and adrenal glands, compared to the broader diencephalon. This may reflect the heightened regulatory role of hormones in these organs, where transcriptional programs are more tightly linked to genotype-dependent hormonal signaling.

To quantify variability of individual gene expression across the eight genotypes, we applied a genotype specificity index (GSI), a metric similar to a tissue specificity index ^104,105^ (Methods). A GSI value of 1 indicates a gene expressed exclusively in a single genotype, while a value of 0 indicates equal expression across all genotypes. After filtering for expressed genes in each cell type of the diencephalon, pituitary gland, and adrenal gland, we examined the proportion of genotype-specific gene expression, defined as genes with a GSI value greater than 0.5. This revealed a broad range of genotype-specific gene expression, independent of the number of nuclei in a given cell type (Fig. 4b). On average, cell types in the pituitary and adrenal glands exhibited a higher proportion of genes that varied significantly across genotypes. We further examined GSI distributions across different regulatory gene classes (Fig. 4c). The results suggest that transcription factors and transcriptional regulators exhibit greater variability between genotypes compared to chromatin state-associated genes and histone remodelers, as determined by a Kolmogorov-Smirnov test against non-regulatory genes. Additionally, clustering genes with a GSI value > 0.5 in pituitary gland cell types showed that genotype-specific genes cluster by cell type rather than genotype (Fig. 4d).

We calculated DEGs in each pituitary cell type, comparing B6 to each of the other seven genotypes. CAST and PWK exhibited the highest number of DEGs across nearly all pituitary cell types, with particularly high variation in lactotrophs and somatotrophs (Fig. 4e). Given their roles in growth and metabolism, these differences may reflect strain-specific variation in endocrine regulation. In contrast, melanotrophs showed a distinct pattern, with 129S1J exhibiting a markedly higher number of DEGs compared to other lab strains. This suggests genetic differences in the regulation of melanocyte-stimulating hormone and its downstream effects on pigmentation. Interestingly, AJ and NOD, which are both albino strains have fewer DEGs in melanotrophs compared to B6 than other strains, other than NZO. Beyond global differences in DEG counts, key pituitary signaling genes also showed genotype-dependent expression variation. For example, *Avpr1b* is a vasopressin receptor that stimulates ACTH release in corticotrophs ^106^ and was predominantly expressed in 129S1, NZO, and the wild-derived strains (Fig. 4f). Meanwhile, *Prlr*, the prolactin receptor, exhibited notably low expression in B6 and somewhat reduced expression in CAST corticotrophs compared to other genotypes (Fig. 4g). Together, these findings suggest that genotype influences both broad transcriptional variation and the expression of critical regulatory genes in the pituitary gland, shaping hormonal signaling and endocrine function differences across mouse strains.

### Satellite cell activation in AJ skeletal muscle

Skeletal muscle stem cells, or satellite cells, were the only cell type where the A/J laboratory strain, (“AJ”), rather than PWK or CAST, had the highest number of DEGs (Fig. 5a, Supp. Fig. 8a). Satellite cells, a rare but essential stem cell population required for adult muscle repair, comprised only 1% of all nuclei in our gastrocnemius dataset. Upon myofiber damage, satellite cells become activated and proliferative. Cycling myoblasts begin to downregulate *Pax7* while upregulating *Myod1* and *Myf5* ^107^. Through asymmetric division, a fraction of *Pax7*+ satellite cells remains quiescent to sustain the stem cell pool ^108^, though *Myf5* expression has also been shown to be maintained in quiescent satellite cells in adult tissue ^109,110^. During myogenic differentiation, *Pax7* expression is fully repressed while *Myod1* expression persists and *Myog* (myogenin) is upregulated. *Myog* continues to be expressed in myotubes along with *Myh3*, a myosin heavy chain isoform known as a marker for newly formed myotubes ^111^, and *Myf6* (MRF4), which remains expressed in mature myofibers where it contributes to fiber maintenance ^112^. Together, the myogenic regulatory factors (MRFs) *Myf5, Myod1*, and *Myf6* exhibit overlapping but partially redundant roles in myogenesis, while *Myog* is essential for myogenic differentiation during development ^107^. Interestingly, we found *Myog* was upregulated in AJ satellite cells compared to 129S1 and WSB (adj. p-value < 0.05 and LFC >1), as well as *Myf6* when compared to 129S1 (adj. p-value < 0.05 and LFC >1). Conversely, the quiescence marker *Pax7* was significantly downregulated in AJ compared to all other genotypes (adj. p-value < 10^−5^ and LFC >1) in addition to *Myf5* (adj. p-value < 0.05 and LFC >1, except compared to PWK). Additionally, *Myh3* was upregulated in AJ compared to every other genotype (adj. p-value < 10^−5^ and LFC >3). Gene ontology and Reactome term enrichment analysis of genes upregulated in AJ highlights terms related to muscle contraction and development, whereas genes downregulated in AJ are enriched for extracellular matrix organization and collagen synthesis (Fig. 5b).

AJ mice are predisposed to late-onset muscular dystrophy due to a dysferlin deficiency caused by a 6,000 bp retrotransposon insertion in intron 4 of *Dysf*, which disrupts splicing ^38,113^. Dysferlin, a sarcolemmal protein, is associated with limb-girdle muscular dystrophy 2B, Miyoshi myopathy, and distal anterior myopathy when mutated ^113^. By 14 months, AJ muscle exhibits inflammation and fatty infiltration comparable to other muscular dystrophy models at 9 months ^114^. Although we do not observe upregulation of inflammatory pathways in AJ mature myonuclei at 10 weeks, several other genes critical for muscle function including *Ryr1, Tnnt3, Mybpc2*, and *Atp2a1* are significantly downregulated in AJ compared to B6J (adj. p-value < 0.01, log2 fold change < −1), suggesting that functional impairments may precede overt inflammatory responses in the progression of muscular dystrophy. In humans, dysferlin is upregulated in activated satellite cells of dystrophic muscle, despite an overall deficiency due to the mutation ^115^. Consistent with this, we detect *Dysf* upregulation in AJ satellite cells but downregulation in mature myonuclei compared to other genotypes (Supp. Fig. 8b). The upregulation of specific MRFs and enrichment of muscle-related GO terms suggest that satellite cells in 10-week-old AJ mice exhibit heightened activation toward myogenesis, potentially reflecting a compensatory response to their predisposition to muscular dystrophy.

To investigate the regulatory programs underlying satellite cell states, we applied topic modeling using Latent Dirichlet Allocation (LDA) with a curated vocabulary of transcription factors, chromatin regulators, and microRNAs ^116,117^. This approach, implemented through the Topyfic package ^116^, identifies latent regulatory topics, which are sets of co-expressed genes that define cellular programs. Each cell is assigned a participation score for each topic, reflecting the extent to which its gene expression profile aligns with that regulatory program. Unlike traditional marker genebased classification, topic modeling captures the complexity of cellular regulation by allowing cells to participate in multiple overlapping programs, providing a nuanced view of transcriptional dynamics across genotypes. To investigate the early impact of dysferlinopathy in satellite cells, we identified nine regulatory topics with varying enrichment across genotypes (Fig. 5c). AJ satellite cells showed the strongest biases, where cell participation is enriched in topics 2, 7, 8, and 9, while topics 1, 3, 4, 5, and 6 were depleted (Fig. 5d, Supp. Fig. 8c). These AJ-enriched topics were also present in cells from other genotypes but at lower levels. For example, topic 2 had the highest representation in AJ, but was also detected in 8.8% of cells across other genotypes. Similarly, topic 8 comprised 11% of AJ satellite cell participation compared to an average of 1.3% in other genotypes, except WSB (6.5%). Topic 7 accounted for 17% of AJ cell participation versus 5.6% in other strains. Structure plots revealed a gradient of participation in AJ-enriched topics across genotypes, where most cells exhibited low to intermediate participation, while a subset was predominantly defined by these topics (Fig. 5d).

Analyzing gene weights associated with these topics identified *Pax7* as highly specific to AJ-depleted topics 4, 5, and 1, while *Mef2c* and *Mir133a-1* are strongly associated with AJ-enriched topics 2 and 7 (Supp Fig. 8d). *Mef2c* synergizes with MRFs to activate myogenesis, while *Mir133a* is a well-characterized “myomiR,” a microRNA essential for skeletal muscle development ^118,119^. The presence of AJ-enriched topics in a subset of cells from other genotypes, albeit at lower levels, suggests that this myogenic program is not inherently misregulated but is more frequently activated in AJ satellite cells compared to other genotypes. To refine our analysis, we subclustered satellite cells based on their participation in regulatory topics. Cells are grouped by high participation in specific topic combinations by hierarchical clustering: subcluster 1 (topics 4 and 5), subcluster 2 (topics 1, 3, and 6), subcluster 3 (topic 9), and subcluster 4 (topics 7, 2, and 8) (Methods, Fig. 5c, e). Marker gene expression profiles distinguish these clusters as quiescent *Pax7*+ satellite cells (subcluster 1), a *Pax7*-low intermediate state (subcluster 2), activated satellite cells (subcluster 3), and *Myog*+ myoblasts (subcluster 4) (Fig. 5f). At this resolution, expression patterns in AJ satellite cells largely align with those of other genotypes. For example, *Pax7*+ AJ satellite cells closely resemble *Pax7*+ satellite cells from other genotypes, and similar trends are observed across other subclusters. However, notable differences emerged, such as significantly higher *Myh3* expression in subclusters 3 and 4 of AJ compared to other genotypes. Overall, these findings suggest that AJ satellite cells are not fundamentally dysregulated at the transcriptional level. Instead, a greater proportion of AJ satellite cells are activated towards myogenic differentiation, with a corresponding reduction in the fraction of self-renewing stem cells.

A study conducted on satellite cells isolated from dystrophic mouse muscle demonstrated that their regenerative capacity remained intact compared to controls, implying that the *in vivo* environment plays a crucial role in regulating satellite cell function ^120^. To investigate the impact of myogenic over-activation on satellite cell signaling in AJ mice, we inferred intercellular communication between cell types based on ligand-receptor interactions within each genotype ^121^ (Methods). We quantified the number of interactions between cell types, including satellite cell subclusters. Overall, *Pax7*+ quiescent satellite cells exhibited 1.6-fold fewer ligand-receptor interactions with other cell types compared to *Myog*+ myoblasts, with 407 and 685 total unique interactions, respectively, across all genotypes (Fig. 5g). AJ myoblasts displayed the highest number of autocrine interactions (subcluster 4 signaling to itself), suggesting a highly dynamic population. As signaling targets, AJ myoblasts exhibit the highest number of interactions with almost every other cell type, with fibro-adipogenic progenitor cells, adipocytes, and activated satellite cells (subcluster 3) being the most prominent sources (Fig. 5g). This extensive signaling within the satellite cell population is likely biologically significant, as the mechanisms governing asymmetric division and maintenance of the stem cell pool in muscle remain incompletely understood. Among the AJ-specific ligand-receptor interactions in subcluster 4, several feature *Itgb1* (integrin beta-1) as the receptor, with *Vegfa, Timp2, Tgm2, Nampt*, and *Angptl2* as ligands. *Itgb1* is upregulated in AJ compared to B6 (adj. p-value < 0.01, LFC > 1) and its signaling has been implicated in muscular dystrophy ^122^. Additionally, we identified interactions where *Cd44* serves as the receptor for signals from *Vegfa, Pkm, Col6a1, Lpl, Lama5, Timp2*/*3, Hbegf*, and *Fgf1. Cd44* is upregulated in AJ satellite cells compared to every genotype except NZO and is known to regulate myoblast migration and differentiation ^123^. Furthermore, *Bmp4* signaling from subcluster 1 to *Acvr2b*/*Bmpr1b* in subclusters 3 and 4 was detected, supporting the role of BMP signaling in muscle regeneration ^124^. Interestingly, we also detected *Tnfsf10*-*Ripk1* signaling uniquely in AJ, where subcluster 4 signals to itself and subcluster 3. *Tnfsf10* is also upregulated in AJ compared to every genotype except PWK. Given that *Tnfsf10* (TRAIL) and *Ripk1* are involved in autophagy ^125^, this interaction may represent an early molecular signature of dystrophic muscle turnover or degeneration in AJ. In summary, our findings reveal the over-activation of myogenic regulatory programs in AJ satellite cells compared to other genotypes, potentially contributing to its predisposition for earlyonset dysferlinopathy. While AJ mice are widely used across many research fields, including cancer and emphysema ^37,126^, researchers should consider the full spectrum of phenotypes inherent to this genotype and strain. Our results demonstrate that genotype and strain-specific functional changes in gene expression emerge within this critical cell type as early as two months of age.

## Discussion

The 8-cube founder dataset of 5.2 million nuclei across 516 samples represents a comprehensive catalog of transcriptional variation across genetically distinct strains and both sexes that can be used to explore the impact of genomic variation on gene expression. The impact of strain background is pertinent to research using mouse models, in which genetic background has often been overlooked or not fully reported despite its potential to produce a profound impact on experimental outcomes ^127,128^. By systematically characterizing the transcriptional landscape across multiple genotypes and sexes, we demonstrate significant variability in transcription between different strains that highlights the importance of considering mouse strain and genetic composition in experimental design and interpretation. Our findings challenge a common implicit assumption that C57BL/6J (B6) mice in some way represent a prototypical mouse strain. Instead, when examined at single-cell resolution, B6 exhibits levels of transcriptional divergence similar to other laboratory strains, reinforcing the idea that it is not uniquely representative. By providing a coordinated dataset encompassing both males and females across various strains and subspecies, the 8-cube dataset shows the impact of sex on gene expression, per cell type, in various genetic contexts. And because the Collaborative Cross panel is derived from these strains, the 8-cube resource provides a framework for identifying and contextualizing genotype-dependent transcriptional variation. Beyond this, the dataset can be mined to generate hypotheses linking known physiological or pathological traits in the founders to cell-type-specific gene expression. For instance, our analysis of AJ satellite cells revealed transcriptional signatures consistent with early activation and dysregulation, aligning with AJ’s known susceptibility to late-onset muscular dystrophy. Similarly, the expansion of Kupffer cells in NZO livers corresponds to this strain’s predisposition to obesity and insulin resistance. These and other examples illustrate how the 8-cube data can be leveraged to connect molecular profiles with phenotypic traits, enabling both retrospective interpretation and hypothesis generation. This study also underscores the significance of sex as a biological variable and its impact on gene expression. While sex differences in certain tissues have been well-documented, our analysis across diverse genotypes reveals additional sex-biased transcriptional patterns that extend beyond previously characterized tissues.

While our data offers evidence of transcriptome differences as a function of genotype and sex, some care should be taken in their interpretation and use. One notable challenge is the possibility of inadvertent association between estrus stage and genotype. Estrus stage was determined but for technical reasons could not be made uniform. For example, we observed that our PWK females were predominately in estrus, whereas CAST were in diestrus or proestrus. This uneven distribution of estrus stages in different genotypes confounds the differentiation of genotype-specific effects from those attributable to estrus stage, especially in tissues where estrus stage has significant impact, such as the ovary or reproductive tract. While females were housed in individually ventilated cages, earlier exposure to male pheromones during shipment or shortly before or after arrival could have triggered the Whitten effect, a well-known phenomenon in which grouped females synchronize their estrus cycles following exposure to male-derived pheromones ^129,130^. Additionally, the timing of tissue collection is a recorded variable to consider. Samples were collected sequentially in the same order and time-frame for all strains, except that NZOs were initiated two hours later. (Supp. Table 1). From a data processing standpoint, all data were aligned to the standard GRCm39 reference genome, which is derived from C57BL6/J. This allowed for consistent comparisons across samples but may introduce alignment bias, particularly for the more genetically divergent wild-derived strains. While these biases are unlikely to affect the overall trends, they could contribute to strain-specific underrepresentation or misalignment of certain transcripts. With the recent release of the first telomere-to-telomere (T2T) genomes ^131,132^, future analyses of the eight founders and CC strains using pangenome references will help further refine our understanding of genotype-specific expression.

Our findings highlight known and novel genotype-differential RNA signatures that map to pertinent cell types and functions, even in young adult mice where overt physiological symptoms are not yet apparent. In some instances these presage later overt physiological symptoms. Observations of this kind depended on single-nucleus resolution, tissue composition, number of nuclei captured per tissue, and depth of sequence. Profiling millions of nuclei across multiple coordinated tissues captured cell type interactions within individual tissues but also showed the ability to uncover responses across different tissue systems, as illustrated by the HPA axis. The scale of this dataset enabled detection of strain differences in some relatively minor cell types and cell states easily obscured in bulk RNA-seq approaches. Satellite cells comprise less than 1% of the muscle dataset, highlighting the enhanced granularity afforded by single-cell analysis. Importantly, the resolution of the analysis shapes the ability to detect genotype effects. Although genotype accounts for only 1% of the total variation at the global level(Fig. 1e), its impact becomes much more pronounced when tissues are analyzed at the level of individual cell types. By providing a deeply annotated single-nucleus reference across the eight founder strains, we establish a framework for uncovering transcriptional signatures linked to genotype, sex, and cell type across diverse tissues. We expect it to be valuable for researchers studying strain-specific biology, interpreting results from CC lines, and generating testable hypotheses that bridge genomic variation to phenotypic diversity.

## Data and code availability

### Data availability

IGVF measurement sets (raw fastqs) and intermediate analysis sets (counts matrices) are listed in Supp. Table 2; final analysis objects (AnnData h5ad files) are listed in Supp. Table 4.

### Data processing pipeline

https://github.com/mortazavilab/parse_pipeline

### Cell type annotation and figure generation code

https://github.com/mortazavilab/8cube_manuscript

## Acknowledgements

We thank the UCI Transgenic Mouse Facility for housing the mice and use of their facilities and UCI GRTH for sequencing the libraries. The TMF and GRTH are shared resources of the Chao Family Comprehensive Cancer Center, supported in part by the National Cancer Institute of the National Institutes of Health under award number P30CA062203. A.M. and B.J.W. were supported by UM1HG012077. We also thank Dr. Tallie Z. Baram for her expert guidance and insightful feedback on the neuroanatomical context of our diencephalon and pituitary data as well as Dr. Andy McMahon for insightful feedback on kidney annotation.

## Author contributions

A.M., B.J.W., L.P, and G.R.M. designed the project. S.K. acquired mice and oversaw their husbandry. S.K., G.R.M., and B.A.W. dissected tissues. E.R., H.Y.L., P.M., M.D., R.M., E.T., N.F., N.M., H.Z. and G.F. helped with sample collection as well as performed nuclei isolation, barcoding, and library preparation. A.S.B., M.C., R.K.L., F.R., and I.B.H. provided input on the experimental design. D.K.S. developed the genotype demultiplexing package. D.T. uploaded data and metadata to the IGVF portal. F.R. provided significant input and code for data preprocessing workflow. E.R., R.W., E.A., and N.S. performed data analysis, generated figures, and wrote and edited the manuscript with significant input from A.M., B.J.W., L.P., and G.R.M.

## Methods

### Mice and tissue collection

Mice were obtained from The Jackson Laboratory (Bar Harbor, ME) and housed at the UCI Transgenic Mouse Facility under controlled conditions. All animal procedures were approved by the UCI Institutional Animal Care and Use Committee. Metadata for each animal and tissue, including mouse ID, sex, date of birth and euthanasia, time of euthanasia, dissector ID, body and tissue weights, and estrus stage, and zeitgeber time (ZT), defined as the number of hours since lights on at 06:30 (ZT0), are detailed in Supp. Table 1.

Animals were housed in autoclaved individually ventilated cages (SuperMouse 750, Lab Products, Seaford, DE) containing autoclaved corncob bedding (Envigo 7092BK 1/8” Teklad, Placentia, CA) and two autoclaved 2” square cotton nestlets (Ancare, Bellmore, NY) plus a LifeSpan multi-level environmental enrichment platform. Tap water (acidified to pH2.5-3.0 with HCl then autoclaved) and food (LabDiet Mouse Irr 6F; LabDiet, St. Louis, MO) were provided *ad libitum*. Cages were changed every 2 weeks with a maximum of 5 adult animals per cage. Room temperature was maintained at 72 ± 2°F, with ambient room humidity (average 40-60% RH, range 10-70%). Light cycle was 14h light / 10h dark, lights on at 06:30h and off at 20:30h.

Euthanasia was performed by anesthesia using isoflurane, followed by decapitation for the collection of blood in EDTA-coated tubes (BD cat. #367856). A pap smear was conducted on female mice before tissue collection and stored on Superfrost slides (Fisher Scientific cat. #12-550-15). Organs and tissues were dissected by three expert dissectors working in parallel: brain regions (left and right cortex and hippocampus, cerebellum, and diencephalon and pituitary), trunk organs (heart, lungs, liver, adrenal gland, kidney, gonads, perigonadal fat, and brown adipose tissue), and specific hindlimb muscles (soleus, plantaris, gastrocnemius, tibialis anterior, and EDL). Trunk and muscle tissues were washed in ice-cold HBSS. Tissues were flash-frozen in liquid nitrogen and biobanked at −135°C until further processing. Estrus stage was determined by crystal violet staining of cells from a pap smear with observation by light microscope.

### Isolation of nuclei from mouse tissues

Eight replicates (four male replicates and four female replicates) of each tissue from each of the founder genotypes were processed per day. Flash-frozen tissues were transferred to a chilled gentleMACS C Tube (Miltenyi Biotec cat. #130-093-237) with Nuclei Extraction Buffer (Miltenyi Biotec cat. #130-128-024) supplemented with 0.2 U/ul RNase Inhibitor (NEB cat. #M0314L) on ice. Nuclei were dissociated from whole tissues using a gentleMACS Octo Dissociator (Miltenyi Biotec cat. #130-095-937). The suspensions were sequentially filtered through 70 um and 30 um strainers (Miltenyi Biotec cat. #130-110-916 and #130-098-458, respectively). Nuclei were then resuspended in cold PBS (Life Technologies cat. #15260037) with 0.1% BSA (Life Technologies cat. #15260037) and 0.2 U/ul RNase inhibitor for manual counting using a hemocytometer and DAPI stain (Thermo cat. #R37606). Debris removal solution (Miltenyi Biotec cat. #130-109-398) was applied to gastrocnemius tissue, forming a density gradient to separate nuclei bands from debris layers. For most tissues, 4 million nuclei per sample were fixed using Parse Biosciences’ Nuclei Fixation Kit v2 according to the manufacturer’s protocol. For smaller tissues such as adrenal gland and female gonads, at least 1 million nuclei were used as input for fixation. Briefly, nuclei were incubated in fixation solution for 10 minutes on ice, followed by permeabilization for 3 minutes on ice. The reaction was quenched, then nuclei were centrifuged and resuspended in 300 uL Nuclei Buffer for a final count. DMSO was added before freezing fixed nuclei at −80°C.

### Single-nucleus RNA-seq experiments

Nuclei were barcoded using Parse Biosciences’ Evercode WT Mega Kit v2, following the manufacturer’s protocol. Fixed, frozen nuclei were thawed in a 37°C water bath and added to the Round 1 reverse transcription barcoding plate at 37,500 nuclei per well. The plate design alternated females and males across columns. A majority of the plate comprised a main tissue, where each individual sample was loaded into a unique well (64 wells in total for the 8 genotypes and 4 male and female replicates). The remaining 32 wells contained a multiplexed tissue, different from the main tissue, where two replicates were pooled from two distinct genotypes in the same well. The first barcoding step consists of *in situ* reverse transcription and annealing of the first barcodes, which use a mix of oligo-dT and random hexamer primers to capture full-length mRNA ^52,133^. The nuclei were then pooled and distributed into 96 wells of the Round 2 ligation barcoding plate for *in situ* barcode 2 ligation. Finally, nuclei were pooled again and redistributed into 96 wells of the Round 3 ligation barcoding plate for *in situ* barcode 3 + UMI + Illumina adapter ligation. Finally, nuclei were counted using a hemocytometer and distributed into 15 subpools of 67,000 nuclei per subpool. The nuclei in each subpool were lysed, and cDNA was purified using AMPure XP beads (Beckman Coulter cat. #A63881). The barcoded cDNA then underwent template switching and amplification. After cleaning the cDNA using AMPure XP beads and performing quality checks with an Agilent Bioanalyzer, the libraries were prepared for Illumina sequencing. The cDNA samples were fragmented, size-selected using AMPure XP beads, and Illumina adapters were ligated. To create the final libraries, cDNA fragments underwent another round of amplification, adding the fourth barcode and P5/P7 adapters, followed by size selection and quality check with a Bioanalyzer (Agilent cat. #G2938A). Libraries were sequenced with two runs of the Illumina NovaSeq 6000 sequencer using the S4 Reagent Kit v1.5 300 cycle kit (cat. #20028312) with configuration PE 140/86/6/0. All 15 subpools were sequenced together with 5% PhiX spike-in to an average depth of 20 billion reads per experiment (approximately 20,000 reads per cell).

### Data preprocessing workflow

Cell-by-gene counts matrices quantifying gene expression were generated from NovaSeq fastqs using a Snakemake workflow based on the kallisto bustools suite ^134–136^, https://github.com/mortazavilab/parse_pipeline. We used kallisto bustools to pseudoalign 165,262,030,405 total reads across all experiments and sub-pools to the mm39 genome with GENCODE vM32 annotations, assign reads to genes, demultiplex cell barcodes, and deduplicate UMIs ^135^. We proceeded with the kallisto matrix that includes all mature, nascent, and ambiguous reads (cells_-x_genes.total.mtx) ^136^, where polyA and random hexamer reads coming from the same well are collapsed per nucleus and transcript for consistency with the Parse Trailmaker software ^52^. The counts matrices, gene information, and cell barcodes were compiled into anndata H5ad files ^137^. Auxiliary python code merged detailed sample- and mouse-level metadata into the corresponding observation or “obs” table. Where appropriate, such as in this experimental design, the pipeline uses the genetic demultiplexing package klue (https://github.com/Yenaled/klue) to quantify the number of genotype-specific reads to each multiplexed nucleus. Genome sequences for the mouse strains were downloaded from from NCBI (https://api.ncbi.nlm.nih.gov/datasets/genome/?taxon=10090) to generate a custom reference, and each multiplexed genotype combination (B6 and NOD, AJ and PWK, 129S1 and CAST, WSB and NZO) was compared in each library. Each nucleus was assigned a genotype based on the maximum number of counts in one of the two expected genotypes. In parallel, our workflow runs CellBender ^53^on the raw kallisto counts matrix in order to remove background noise such as highly-expressed tissue-specific transcripts that arose from our experimental design. CellBender parameters were manually adjusted per subpool (Supp. Table 2) to minimize loss of “real” nuclei. Doublet detection was also performed with Scrublet ^138^ using the uncorrected counts for each subpool. Other than a baseline UMI threshold of 200 UMIs per nucleus, no additional filtering was performed. Finally, all nuclei belonging to the same tissue in each experiment were merged into a single H5ad file for downstream tissue-level analysis.

### QC and clustering single-nucleus data

The nuclei were further aggregated at the tissue level after all experiments were quantified so that we could combine nuclei from the same samples that were sequenced on different runs (Fig. 1c). Male and female gonads were processed separately. All tissues were filtered for nuclei with >500 and <150,000 UMIs, >250 expressed genes (>=1 UMI), <1% mitochondrial gene expression, and <0.25 doublet scores. Multiplexed nuclei with ambiguous genotypes (around 0.2% of the dataset) were also excluded from downstream analysis. Scanpy (version 1.10.2) was used to cluster nuclei for cell type annotation ^137^. Briefly, the CellBender-corrected counts matrices for each tissue were normalized by total UMI count followed by logarithmic transformation and filtration for highly variable genes (min_mean=0.0125, max_mean=3, min_disp=0.5). Percent mitochondrial gene expression and number of genes detected were regressed. Dimensionality reduction using PCA with the top 30 principal components was used to construct a neighborhood graph (n_neighbor = 20). Leiden clustering was performed for each tissue at the same resolution (= 1).

### Cell type annotation

Clustered nuclei were manually annotated using expression of known marker genes, discussions with expert collaborators, identification of cluster marker genes and cross-referencing with literature, as well as integration with label transfer from a reference dataset using scVI-tools when possible ^139,140^. Annotation was performed at three levels, from fine to coarse resolution: subtype, cell type, and general cell type. For the most part, each cluster was assigned to a single subtype, with multiple clusters making up larger subtypes. Subclustering and annotation was performed based on clear marker gene expression in certain cell types, such as skeletal muscle myonuclei, adrenal cortical zones, stages of spermatogenesis, loop of Henle regions, and neuroendocrine cells of the pituitary gland. For liver, hepatocytes were annotated and separated from other cell types, which were clustered and annotated independently. Cell types are also annotated by a Cell Ontology (CL) ^141^ ID which matches the “cell type” level annotations. When necessary, “subtypes” extend CL cell types into cell subtypes and/or cell states. For example, “subtypes” captures specialized myonuclei such as those resting underneath the neuromuscular junction and myotendinous junction ^142,143^. The “general cell type” level uses the hierarchical structures embedded in the CL database to group cells into 13 broader annotations such as “epithelial” and “germ” cells (Fig. 2c). In total, we annotated 106 unique cell types and subtypes across all tissues. The number of annotated cell types per tissue was 24 in both diencephalon and pituitary and in cortex and hippocampus, 19 in male gonads, 18 in kidney, 17 in gastrocnemius muscle, 14 in both heart and female gonads, and 12 in both liver and adrenal glands.

### Variance decomposition analysis

For a dataset that can be split into groups, the law of total variance decomposes the variance into two terms: explained and unexplained variance. The explained variance is the variance of the means of each group, and the unexplained variance is the mean of the variances within each group. To generate a list of strain markers, first variance explained by strain and variance explained by cell type were calculated for all genes, in each tissue separately, with male and female gonads treated as separate tissues.

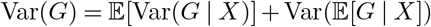

These explained variance terms were used to generate an F statistic, and Benjamini-Hochberg correction with a q-value of 0.1 was applied to extract a list of genes more variable by strain than by cell type in each tissue.

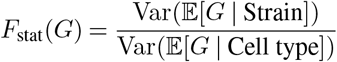

The intersection of the 9 sets of genes was taken. Mean expression values of these genes were observed in all 9 tissues across all 8 strains to identify genes that are strain markers in all tissues.

### Pseudobulk PCA and differential expression analysis

Pseudobulk profiles were generated using CellBender-corrected counts at the level of individual tissue samples (“lab_sample_id”), summing counts per sample using decoupler ^144^ with default parameters (minimum of 10 cells and 1,000 counts). Counts were then normalized to reads per million (RPM). For PCA, counts were additionally log-transformed. Pseudobulk count matrices were merged across tissues, resulting in a final matrix of 40,892 genes by 515 samples. PCA was performed using Scanpy v.1.10.2 on the normalized dataset containing all genes ^137^. Per-tissue PCA (Supp. Fig. 3) was conducted using the same approach but applied separately to each tissue. Differential expression analysis was performed for all 28 pairwise genotype comparisons using PyDESeq2^145^ on non-normalized pseudobulk counts per sample, with sex included as a covariate in non-gonadal tissues. Results were filtered for a maximum adjusted p-value of 0.01 and an absolute log2 fold change greater than 1. Gene ontology and Reactome term enrichment analyses were conducted using GSEApy ^146^ with the Reactome 2022 and GO Biological Process 2023 gene sets. For visualizing gene expression patterns within specific cell types (e.g., violin plots in Fig. 2 f-i, Fig. 3 c-j, Fig. 4 f-g), CellBender-corrected counts were subset for nuclei within the cell type of interest. Counts were then processed using decoupler with default options, followed by RPM scaling.

### Identifying genotype and sex-driven variation through cell type subclustering

For each annotated cell type with more than 50 nuclei in every genotype, we reprocessed the CellBender-corrected counts by normalizing to counts per 10k, log-transforming, and subsetting for highly variable genes using default Scanpy parameters. To focus on cell-type-specific differences, we removed the 82 genotype-specific genes identified using variance decomposition analysis (Fig. 1f, Supp. Table 3). No regression was performed. PCA and nearest-neighbor graph calculations were conducted using default parameters, followed by Leiden clustering with a resolution of 1. Genotype-specific clustering was assessed using a chi-squared test (scipy.stats v1.10.1) ^147^, comparing the observed genotype distribution in each subcluster to the expected distribution within the cell type. Clusters were classified as genotype-specific if the chi-squared p-value was <10^−10^. Genotype-specific clusters where a single replicate accounted for >75% were excluded from the analysis. Nuclei were then categorized based on whether they fell into genotype-specific clusters, enabling visualization of genotype-driven clustering patterns across cell types. A similar approach was applied for sex-specific clustering, except subclustering was performed within each genotype and cell type to minimize genotype-driven effects. The single-replicate filter was not applied in this analysis. Sex-specific clustering was not assessed in gonadal tissues.

### Calculation of genotype specificity index

Gene expression genotype specificity was determined using GSI (genotype specificity index):

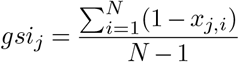

GSI is a modification of the tissue specificity index which has been used for RNA-seq and microRNA-seq data ^104,105^. In this formula, *x*_*j,i*_ represents the expression of gene *j* in genotype *i*, normalized by the maximum expression in any genotype. *N* is the number of genotypes (always 8).

Pseudobulk profiles at the sample level were calculated for individual cell types. GSI calculation was only performed on genes which had greater than 0.5 mean normalized expression in a given cell type. Gene biotypes (Fig. 4c) included transcription factors, chromatin regulators, and microRNA host genes, identified through GO term annotations and microRNA expression correlations ^116,117^.

### Identifying regulatory topics in satellite cells using Topyfic

Topics were calculated using a curated vocabulary of 2,766 regulatory genes with the Topyfic package as previously described ^116^. To ensure reproducibility, Latent Dirichlet Allocation (LDA) was run 100 times with varying initial conditions, and the resulting topics were clustered to identify consensus topics ^116^. Using a resolution of k = 10, we identified 9 robust topics across 8,186 satellite cell nuclei. Structure plots (Fig. 5d) were generated using the structure plot function in Topyfic. Cell participation scores were extracted from the TopModel object for clustering satellite cells into cell states based on topics. Gene weights were also extracted using get_gene_weights to compare the contribution of key genes across topics (Supp. Fig. 8d). Finally, satellite cell subclustering was performed using hierarchical clustering of cell participation scores with a correlation distance metric (scipy v1.10.1) ^147^.

### Ligand-receptor interactions in satellite cells

We analyzed intercellular communication in muscle tissue using LIANA+, a Python package for ligand-receptor interaction inference ^121^. LIANA+ incorporates eight distinct ligand-receptor inference methods, including those from CellPhoneDB ^148^, CellChat ^149^, Connectome ^150^, NATMI ^151^, SingleCellSignalR ^152^, sc-SeqComm ^153^, log fold change, and a geometric mean approach. The skeletal muscle dataset was normalized using counts per 10k and log transformed. For each genotype, we independently filtered the data and input it into LIANA+. We applied LIANA’s rank_aggregate function, which integrates predictions from multiple methods, using the mouse consensus resource. To refine the results, we filtered interactions by CellPhoneDB permutation-based p-values (< 0.05) as a measure of interaction specificity ^148^. Finally, we identified ligand-receptor pairs unique to a single genotype in each cell type interaction in muscle tissue and quantified the number of genotype-specific interactions per cell type (Fig. 5g), highlighting potential strain-dependent differences in signaling networks.

### Supplementary Tables

- **Table S1: Sample collection metadata**.
- **Table S2: CellBender parameters for each plate and subpool as well as IGVF measurement set and intermediate analysis set accessions**.
- **Table S3: Strain marker genes from variance decomposition analysis**.
- **Table S4: IGVF principal analysis set metadata**.

## Supplementary Figures

- **Figure S1: Relationship between body weight and tissue weight in 9 diverse tissues**.
- **Figure S2: Detection and removal of cross-tissue contamination using CellBender**.
- **Figure S3: Pseudobulk principal component analysis in each tissue**.
- **Figure S4: Tissue-specific differential gene expression**.
- **Figure S5: Percent of nuclei subclustering by genotype across cell types and tissues**.
- **Figure S6: Genotype-specific Kupffer cell subclusters and inflammatory signatures**.
- **Figure S7: Subclustering of hypothalamic neurons reveals 39 subclusters**.
- **Figure S8: Differential gene expression and regulatory topic modeling in 8**,**186 satellite cells**.

**Figure S1.**
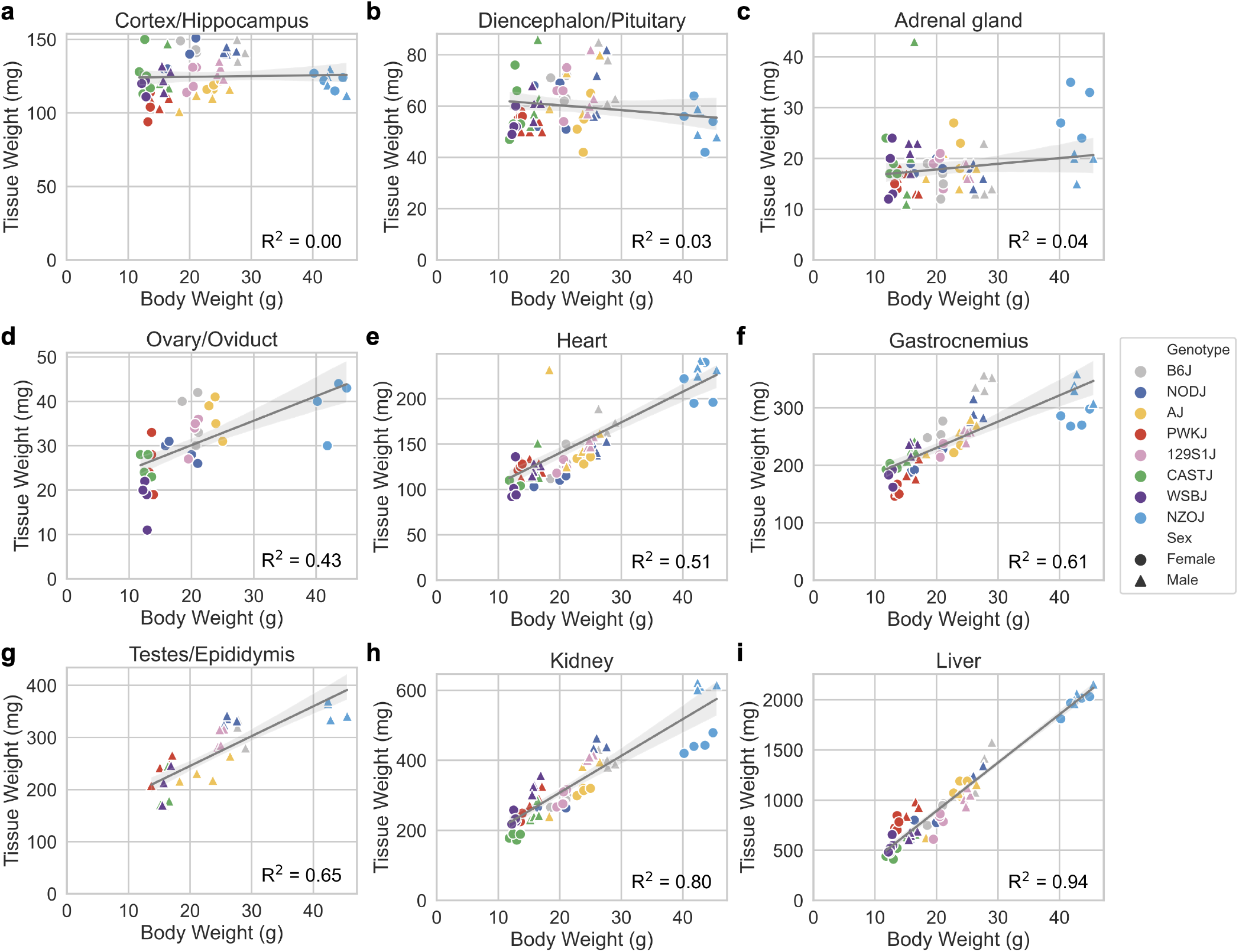
Relationship between body weight and tissue weight in 9 diverse tissues. Body weight compared to weight of each tissue, with linear regression fit (grey line) and Pearson R^2^ in: **a**, Cortex/hippocampus (R^2^ = 0.0) **b**, Diencephalon/pituitary (R^2^ = 0.3), **c**, Adrenal glands (R^2^ = 0.04), **d**, Female gonads (R^2^ = 0.43), **e**, Heart (R^2^ = 0.51), **f**, Muscle (R^2^ = 0.61), **g**, Male gonads (R^2^ = 0.65), **h**, Kidney (R^2^ = 0.80), and **i**, Liver (R^2^ = 0.94).

**Figure S2.**
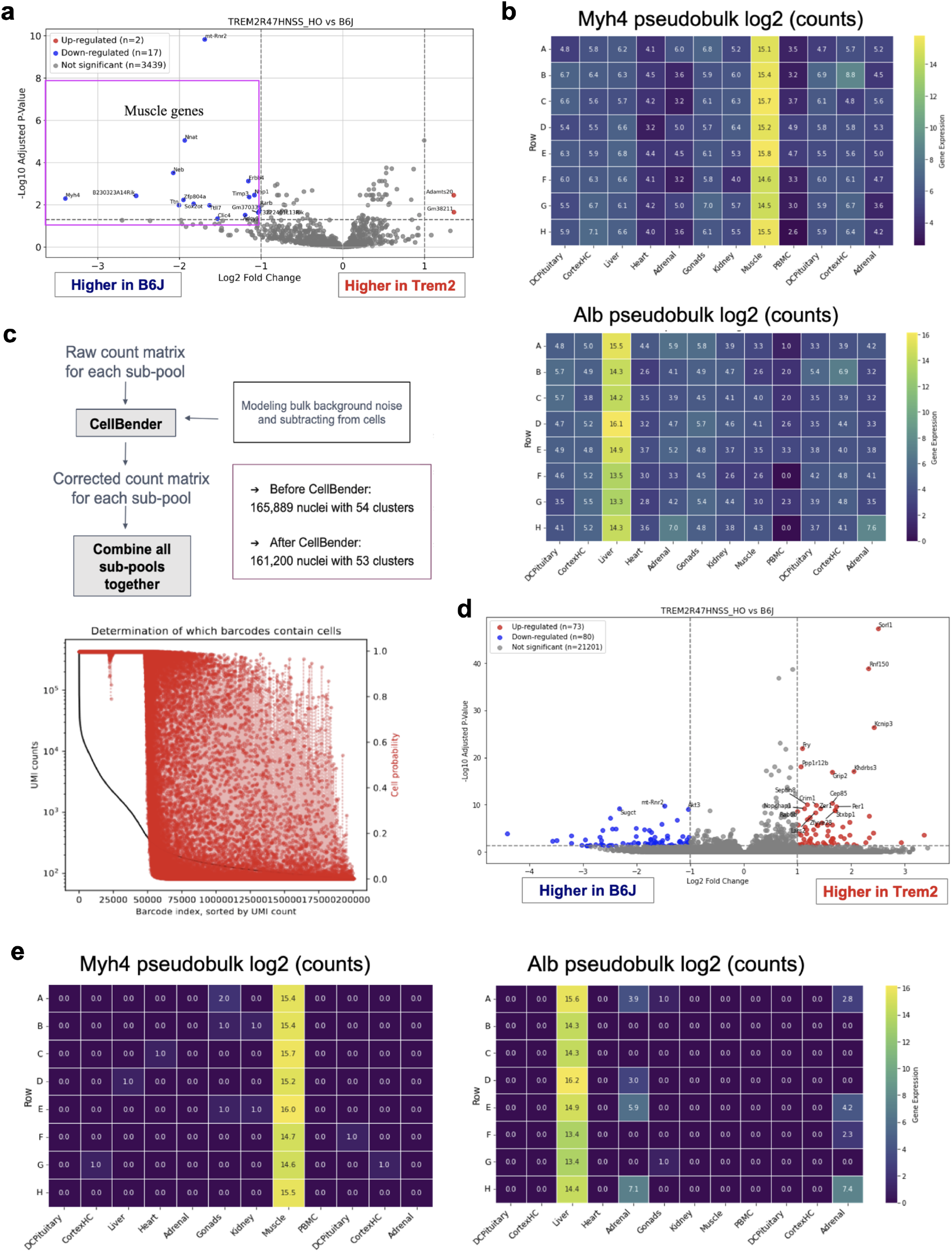
Detection and removal of cross-tissue contamination using CellBender and subsequent pseudo-bulk differential gene expression (DEG) analysis. **a**, Pre-CellBender DEG analysis of brain microglia between *Trem2*^R47H NSS^ homozygotes (“Trem2”) and B6, highlighting contamination from other tissues as indicated by the presence of skeletal muscle (*Myh4*) and liver (*Alb*) markers. **b**, Heatmaps showing pseudo-bulk log2 counts for tissue-specific markers, revealing their presence across unrelated tissues prior to contamination correction. **c**, CellBender correction workflow. Raw count matrices for each subpool were corrected independently, reducing nuclei numbers from 165,889 (54 clusters) to 161,200 (53 clusters), effectively identifying and removing contamination. **d**, Post-CellBender DEG analysis of microglia, showing cleaner tissue-specific gene expression with 73 upregulated (red) and 80 downregulated (blue) genes in Trem2 versus B6. GO Biological Process enrichment reveals upregulated genes associated with cAMP catabolic processes (P=0.0046) and downregulated genes enriched in small molecule metabolic processes (P=0.00054). **e**, Post-CellBender heatmaps for tissue-specific markers demonstrate effective reduction of contamination, with these markers now largely restricted to their expected tissues.

**Figure S3.**
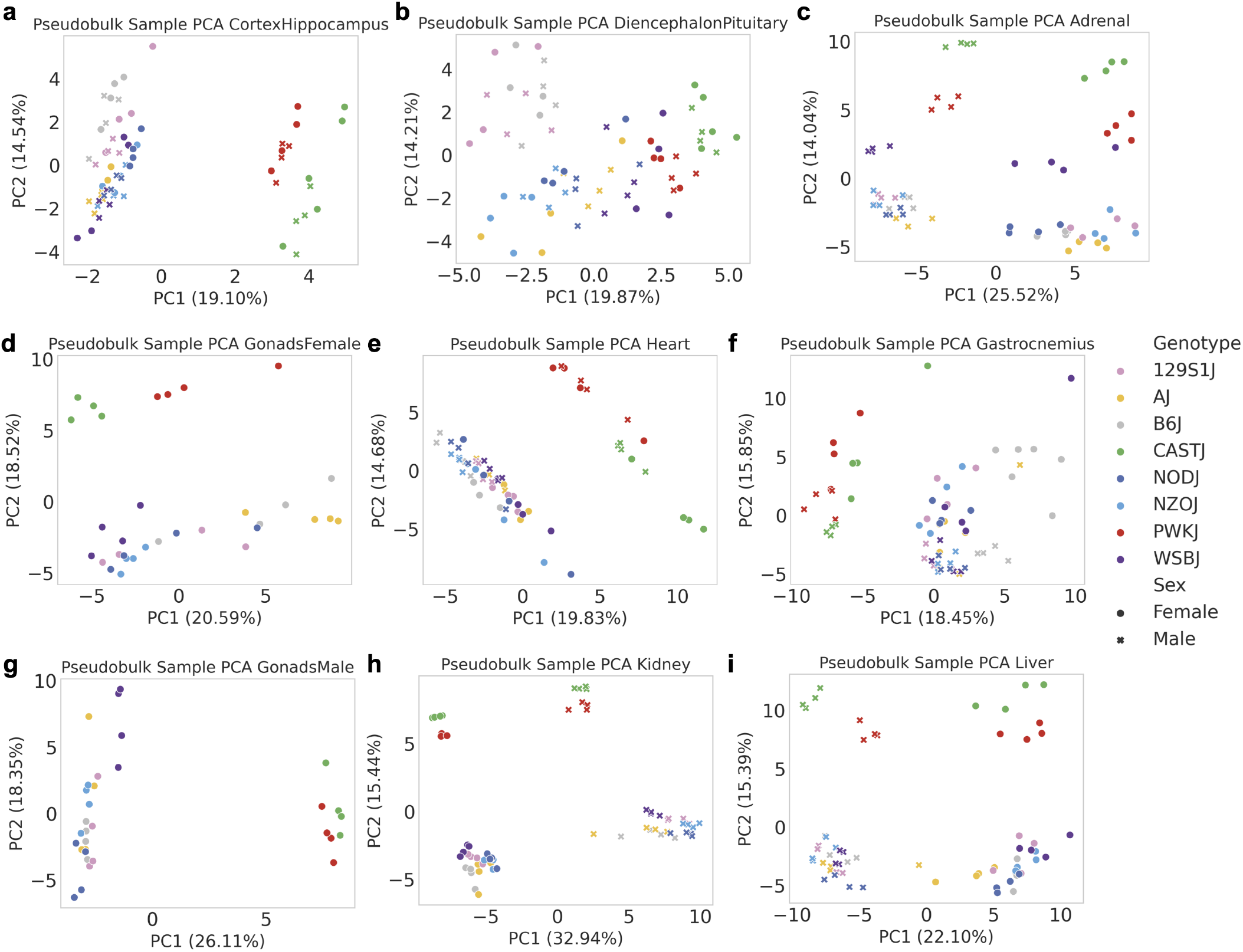
Pseudobulk sample principal component analysis in each tissue. PCA of pseudobulk samples (sum of Cell-Bender counts per sample, followed by read depth and log normalization) in **a**, Cortex and hippocampus, **b**, Diencephalon and pituitary, **c**, Adrenal glands, **d**, Female gonads, **e**, Heart, **f**, Skeletal muscle, **g**, Male gonads, **h**, Kidney, and **i**, Liver.

**Figure S4.**
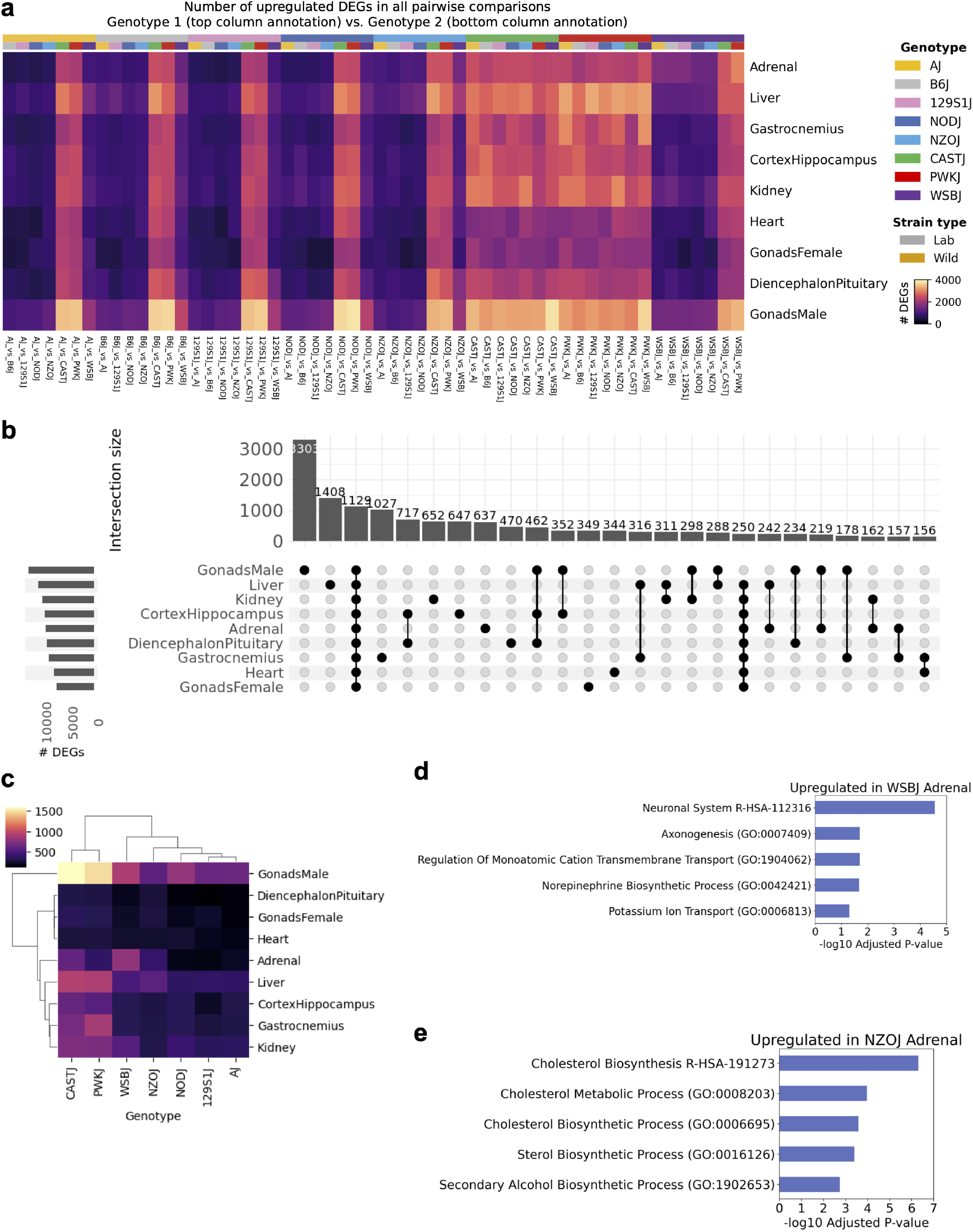
Differential gene expression analysis across tissues. **a**, Number of upregulated DEGs in all pairwise comparisons at pseudobulk sample level across tissues (adj. p-value < 0.01, LFC > 1). The heatmap displays the number of upregulated genes in each genotype (indicated by the top annotation, starting with AJ) when compared to every other genotype (indicated by the second annotation, starting with B6). **b**, Intersections of DEGs from B6 comparisons across tissues. **c**, Number of tissue-specific DEGs in each genotype and tissue. **d**, GO and Reactome terms from adrenal-specific DEGs upregulated in WSB. **e**, GO and Reactome terms from adrenal-specific DEGs upregulated in NZO.

**Figure S5.**
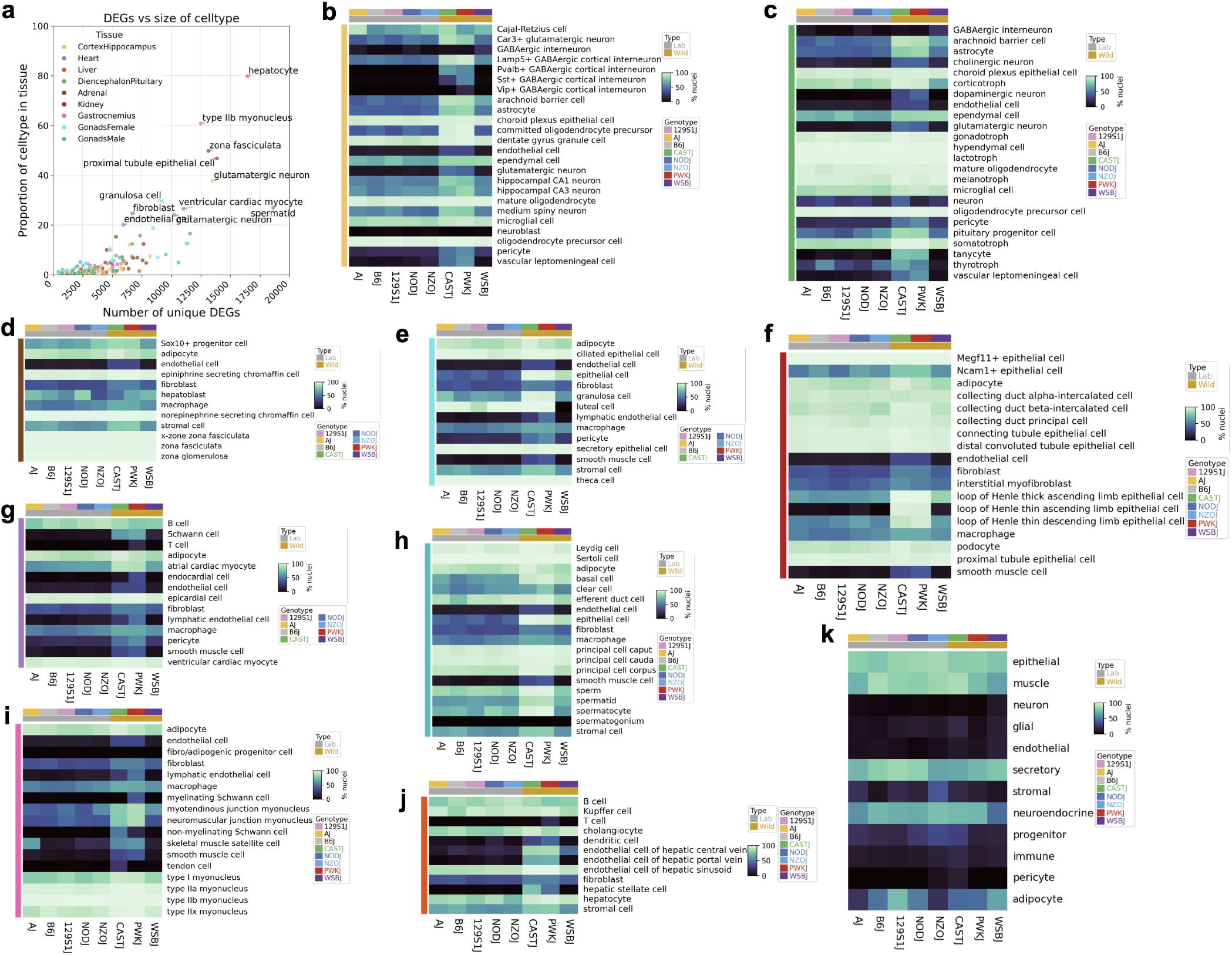
Differential gene expression analysis and subclustering across tissues. **a**, Number of DEGs in B6 comparisons in each cell type compared to the proportion of the cell type in the tissue of origin. b-j, Percent of nuclei clustering by genotype in across cell types and genotypes in **b**, cortex/hippocampus, **c**, diencephalon/pituitary, **d**, adrenal gland, **e**, female gonads, **f**, kidney, **g**, heart, **h**, male gonads, **i**, gastrocnemius, **j**, liver. **k**, Percent of nuclei clustering by sex across general cell types and genotypes.

**Figure S6.**
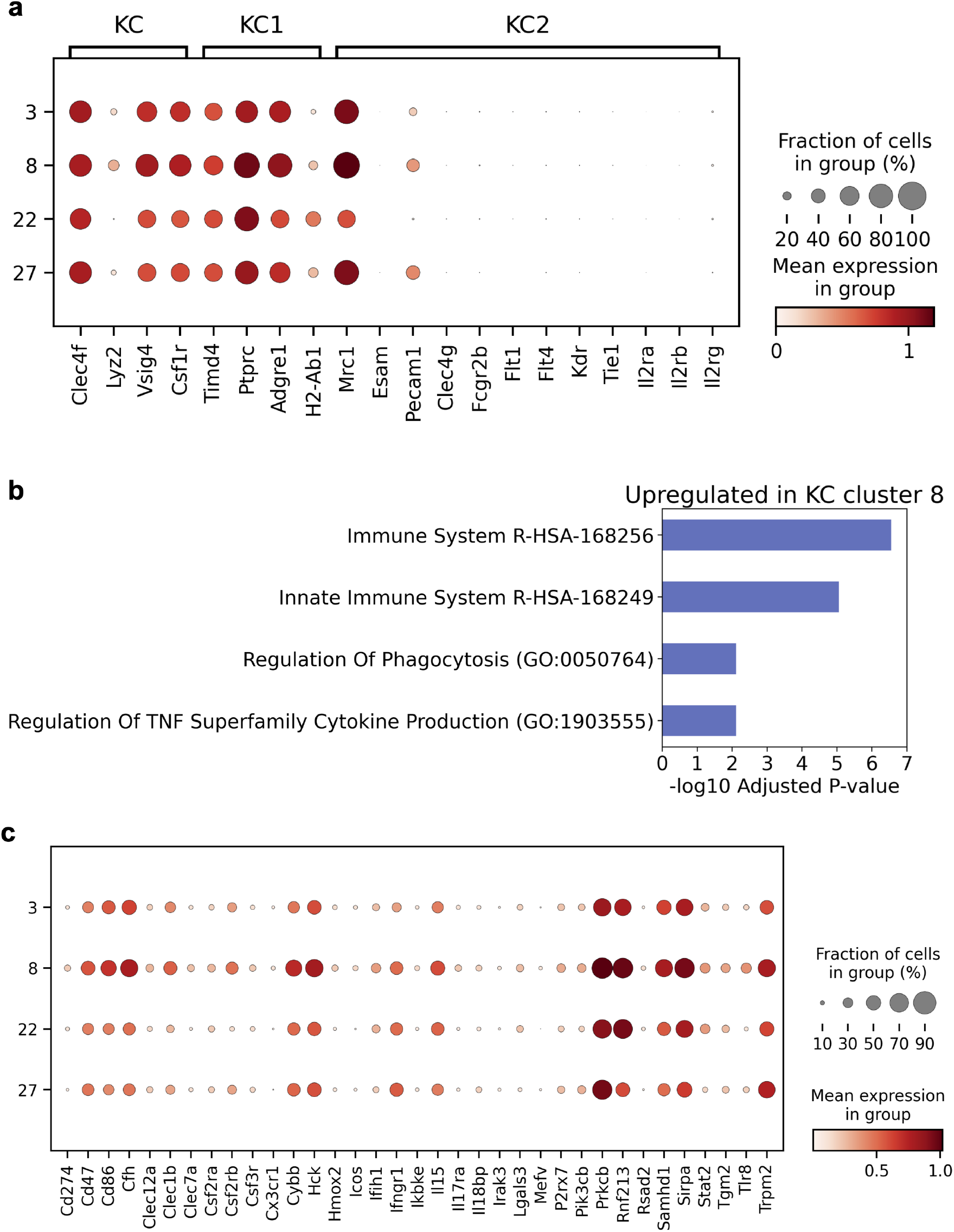
Genotype-specific Kupffer cell subclusters and inflammatory signatures. **a**, Expression of canonical Kupffer cell marker genes as well as KC1 and KC2 markers across immune-focused analysis subclusters ^78,79^. Cluster 8 is NZO-specific, cluster 27 is CAST-specific, cluster 22 is PWK-specific, and cluster 3 is shared across genotypes. **b**, GO term enrichment for genes upregulated in NZO-specific KC cluster 8 (adj. p-value < 0.01). Markers were called using scanpy rank_genes_groups and Wilcoxon rank-sum and filtered for genes upregulated with LFC > 0.5 and adj. p-value < 0.01. **c**, Expression of cluster 8-specific inflammatory marker genes (adj. p-value < 0.01, LFC > 0.5).

**Figure S7.**
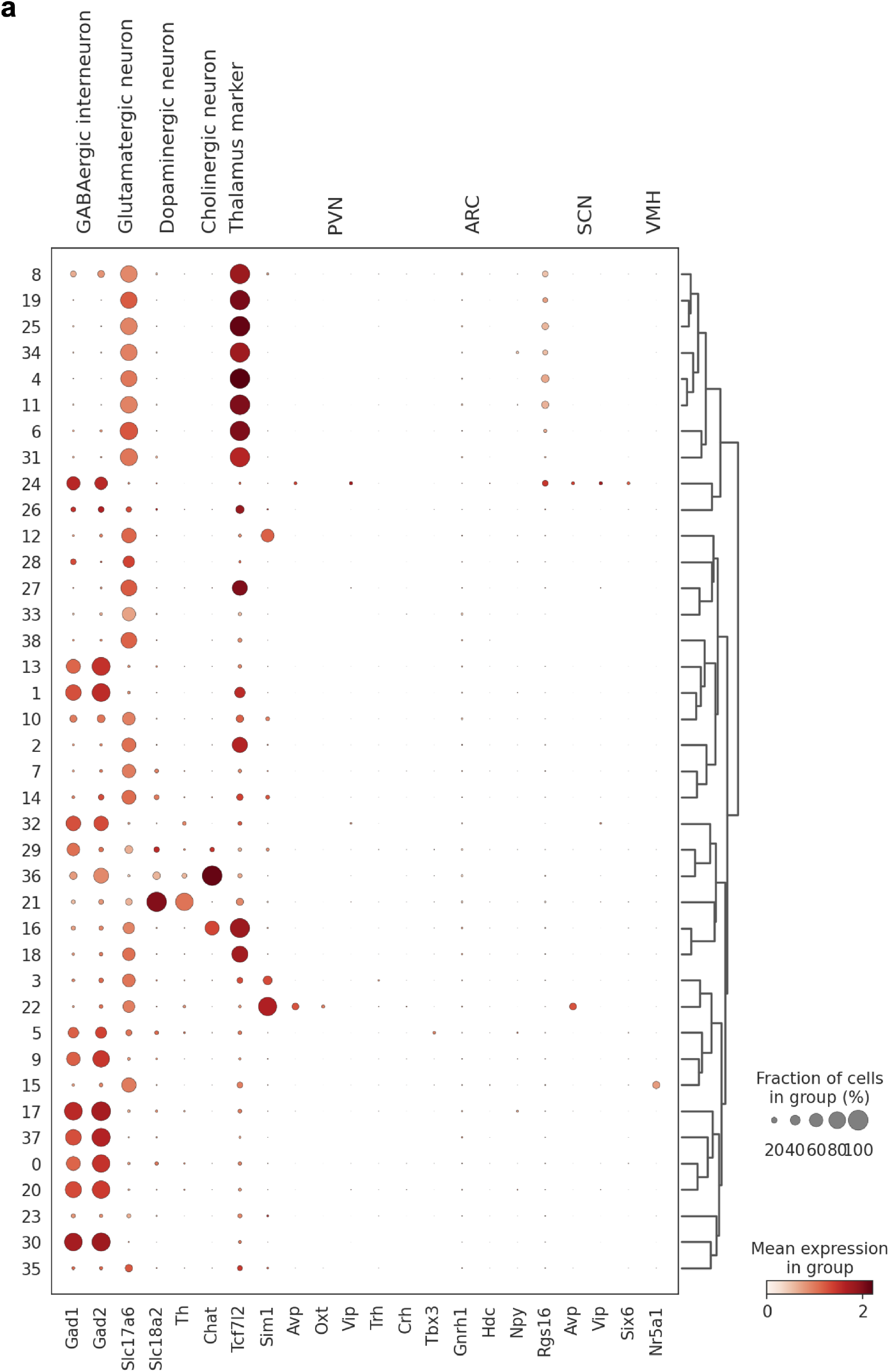
Subclustering of hypothalamic neurons reveals 39 subclusters. **a**, Expression of neuronal marker genes, where each row is a cluster, and each column is a marker gene. Rows clustered by marker gene expression.

**Figure S8.**
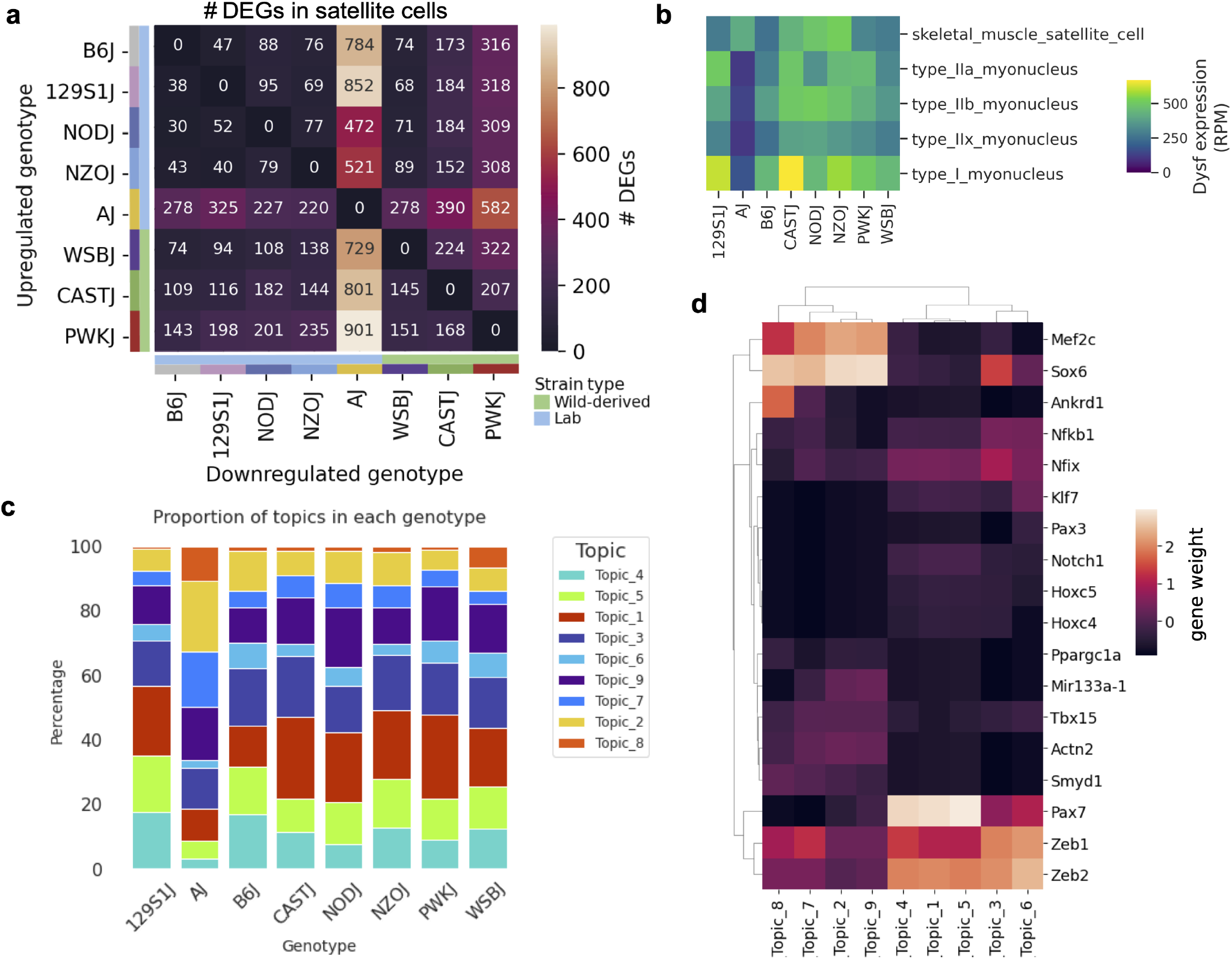
Differential gene expression and regulatory topic modeling in 8,186 satellite cells. **a**, Number of DEGs identified in pairwise comparisons, where the upregulated genotype is indicated in rows and downregulated genotype in columns (adj. p-value < 0.01, |LFC| > 1). **b**, Pseudobulk expression of *Dysf* in skeletal muscle satellite cells and mature myonuclear subtypes across genotypes. **c**, Breakdown of topic participation in nuclei grouped by genotype. **d**, Weight of selected genes across regulatory topics.

